# Pseudo-helicase motifs of SecA involved in ATP-driven dynamic coupling for protein translocation

**DOI:** 10.64898/2026.11.24.768852

**Authors:** Dulcie Havers, William J. Allen, Tom A Williams, Mark S Dillingham, Ian Collinson

## Abstract

The ubiquitous Sec translocon is a highly conserved machine specialised for polypeptide translocation through lipid bilayers. In the bacterial system, the cytoplasmic motor ATPase SecA associates with the core channel complex SecYEG, through which unfolded polypeptides are driven across the membrane by a Brownian ratchet mechanism. How the energy released during the ATP-hydrolytic cycle is transduced to the conformational changes required for preprotein secretion is not well understood. Here, we re-explore the known homology between SecA and the nucleic acid helicases to better understand the fundamentals of dynamic energy coupling during biopolymer transport: how ATP-turnover is regulated by the engaged polypeptide, and how the ensuing conformational changes bring about transport. Using bioinformatics and comparative structural analysis we identify several motifs within SecA equivalent to those required for helicase activity. We then probe their involvement in protein translocation experimentally by site specific amino acid substitution and the analysis of reconstituted variant activity. The results show that the pseudo-helicase translocation motifs are important for regulating ATP-coupled protein transport, enabling the motor to sense and respond to the translocating polypeptide.

## Introduction

The encapsulation and compartmentalisation of cells by phospholipid membranes necessitated the evolution of protein transport machines for extra- and intra-cellular trafficking. One such protein translocon is the ancient protein secretory (Sec) system— essential in every cell in every organism. The bacterial Sec-machinery is responsible for the majority of protein secretion across the plasma membrane, and beyond^1–4^. The protein channel through the membrane is formed by the heterotrimeric SecYEG core-complex^5^, through which unfolded proteins are translocated by the associated SecA, driven by ATP and the PMF^1,6–8^. The molecular motor component SecA is known to associate with the cytoplasmic loops of SecY (**Figure 1A**)^9^.

**Figure 1.**
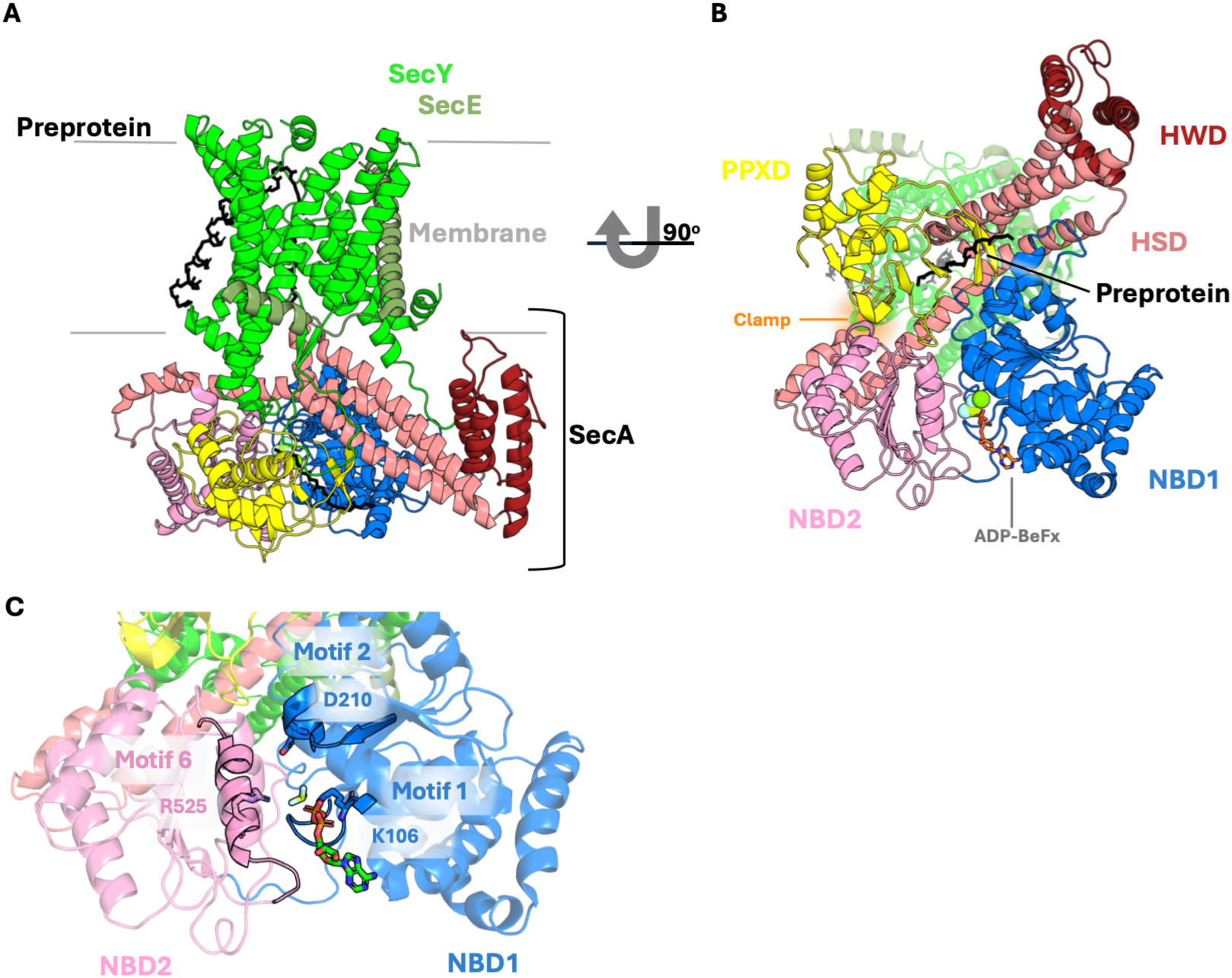
SecA bound to SecYEG, (PDB:7XHA, Dong et al). (A) SecA, shown in multicolour, binds on the cytoplasmic side of SecY, shown in green. Small regions of preprotein have been resolved (shown as black sticks), in both the lateral gate of SecY, and the SecA cavity. SecE is shown in a dark green, the structure was determined in the absence of SecG. (B) SecA contains 5 domains. NBD1 (Blue) and NBD2 (Pink) form the NBS, where ADP-BeFx is shown bound as sticks. Between these domains sits the preprotein cross linking domain (PPXD, yellow). Behind the NBDs sits the helical scaffold domain (HSD, coral) and the helical wing domain (HWD, red). (C) Representation of the three conserved helicase motifs involved in nucleotide binding and hydrolysis. The motifs are shown in bold, with sticks representing key conserved residues. Illustrated using a B. subtilis SecA structure (PDB:7xha) ^C^. Motif 1 - Walker A containing conserved lysine and threonine residues that either directly or indirectly bind the phosphate moieties of ATP. Motif 2 - Walker B, contains conserved aspartate and glutamate residues which, respectively, co-ordinate an essential Mg^2^+ at the active site and act as the catalytic base to activate water for nucleophilic attack at the gamma-phosphate of ATP. Motif C -Arginine finger, containing a conserved Arginine residue involved in stabilising the transition state and polarizing the gamma-phosphate of ATP.

Energy from the ATP hydrolytic cycle of SecA and the trans-membrane proton-motive-force (PMF) is used to drive translocation of proteins with a secretion targeting sequence (preproteins)^10^ through SecYEG^11–13^. SecA is known to undergo large conformational changes during the hydrolytic cycle, and these have been shown to propagate throughout the translocon^6,14–16^. While the dynamic action of SecA is well characterised, it is unclear how the energy from the ATP turnover is transduced to drive the directional movement required for protein translocation^17–20^.

SecA contains five domains (**Figure 1B**), two of which are nucleotide binding domains (NBD1 and NBD2) each displaying a P-loop fold and which together form the nucleotide binding site (NBS) at their interface. The preprotein cross-linking domain (PPXD) is a highly mobile domain that sits close to the interface between the two NBDs. The helical wing domain (HWD) connects to the back of the NBDs and continues into the helical scaffold domain (HSD), a part of which (the two helix finger; 2HF) protrudes into the SecY channel. SecA forms a chamber for preprotein substrate to pass through and responds to nucleotide exchange, with the clamp between the PPXD domain and NBD2 closing when the SecY channel opens upon ATP binding^21^.

Together, NBD1 and NBD2 form a structural unit that closely resembles a nucleic acid helicase core^6,22,23^ (**Figure 1B**). Helicase-like proteins are highly abundant and have been grouped into several superfamilies based on their oligomeric states and primary sequences^23^. Superfamily 1 and 2 enzymes (SF1 and SF2) are monomeric or dimeric proteins containing several conserved ‘signature motifs’, all of which localise to the two core helicase domains (the N- and C-cores). SecA has previously been categorised as a member of SF2^24^, and its NBD1 and NBD2 domains are equivalent to the so-called N-core and C-core domains of helicases, respectively. In helicase-like proteins, these core domains transduce energy from ATP binding and hydrolysis to effect mechanical work on nucleic acid substrates: most commonly unidirectional translocation along single- or double-stranded DNA, which underpins strand separation or remodelling activity^25,26^. The conservation of this helicase core between helicases and SecA highlights a common ancestry with an ancient translocation machinery. Therefore, a level of commonality could exist between the mechanistic features involved in the translocation of protein and nucleic acids. It is tempting to speculate that the helicase motifs play a similar role in chemo-mechanical energy transduction in SecA. We reasoned that an in-depth comparison with the very well characterised coupling between nucleotide turnover and mechanical work in conventional helicases would help us better understand dynamic energy transduction within SecA’s.

The conserved signature motifs of helicases fulfil two main functions: some are directly involved in ATP binding and hydrolysis and located at the interface between the core domains, whereas others are involved in substrate translocation and line the surfaces of the core domains that interact with substrate. The ATPase motifs include Walker A (also called helicase motif 1) and Walker B (helicase motif 2), as well as the arginine finger (helicase motif 6). These are all conserved in SecA (**Figure 1C**), and play well-defined and essential roles in ATP binding and hydrolysis^23^. The translocation motifs, which differ somewhat between the helicase superfamilies, unsurprisingly, are responsible for coupling ATPase activity to nucleic acid translocation *via* an inchworm mechanism^24,27,28^.

In this work, we further explore SecA’s helicase translocation motifs and investigate their potential role in polypeptide transport. Using a structural bioinformatics approach we define ‘pseudo-helicase motifs’ in SecA orthologues and demonstrate experimentally their involvement in the regulation of ATP hydrolysis and polypeptide translocation.

## Results

### SecA is closely related to DEAD-box helicases

To establish the evolutionary relationship between SecA and helicases we first carried out phylogenetic analysis to construct a maximum-likelihood phylogenetic tree using IǪ-Tree2^29^. A database of representative helicase sequences was assembled using a diverse proteomes dataset (See Materials and Methods), that provided representative coverage of the known genomic diversity of prokaryotes. The resulting phylogenetic tree confidently separates SecA from SF1 (**Figure 2A**) (SH-aLRT 90.9%, UFBoot of 64), but the position of SecA within SF2 is unresolved. The dataset was designed to be diverse, so confidence is limited by the gappy alignments that result from this sequence diversity, limiting the resolution of deep relationships in the tree. Consistent with prior research^22,23,30,31^, the fewest base substitutions were between SecA and the DEAH/DEAD-box subfamilies^32^. There is no definitive closest relative for SecA in this unrooted tree, but the closest branches are those of the DEAH/DEAD-box and T1R/T3R subfamily helicases. We next used Foldseek and DALI to perform structural homology searches using *B. subtilis* SecA (PDB:7XHA) as the input. Ignoring SecA orthologues, the top hits using either approach were helicase members of the DEAD-box family (Supplementary Data 1 and Supplementary Table T1), consistent with the close evolutionary relationship

**Figure 2:**
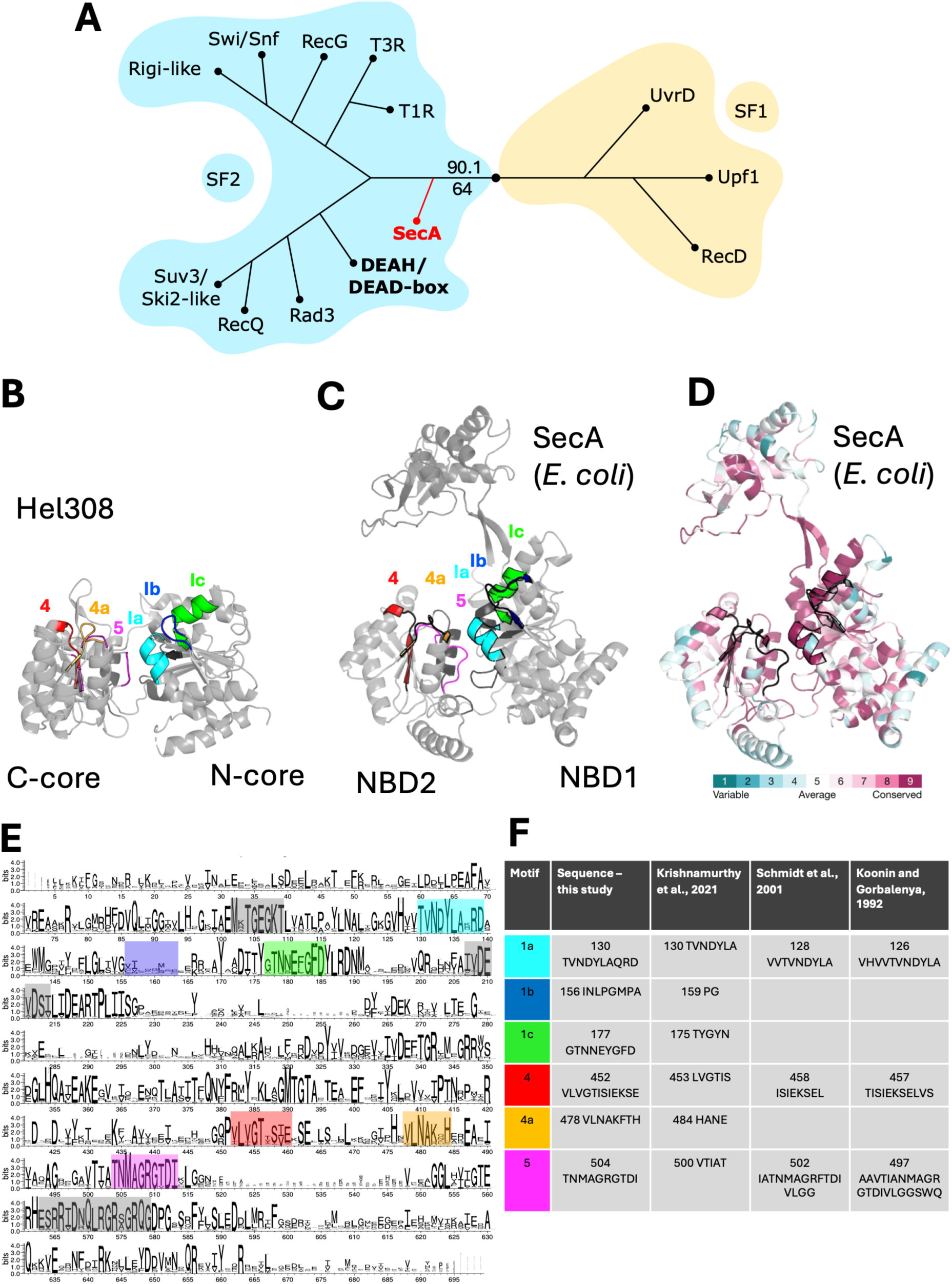
Identification of SecA pseudo-helicase motifs. (A) SecA is most similar to SF2 helicases. SecA is separated from SF1 with good confidence for this tree (SH-aLRT of S0.S and UFBoot of C4). SecA is not grouped alongside any smaller SF2 helicase family, however the results revealed the fewest number of substitutions between SecA and any other helicase family was the DEAH/DEAD box family. (B) Helicase motifs in Hel308 as defined by Büttner et al 2007, with the helicase hydrolysis motifs shown in dark grey, and the helicase translocation motifs shown in bold colours, (PDB:2PCR)^33^. (C) Pseudo-helicase motifs in corresponding structural regions in SecA, (E. coli AlphaFold2 model), again the pseudo-helicase hydrolysis motifs are shown in grey, and the corresponding structural regions of the pseudo-helicase translocation motifs are shown in bold colours, revealing the strikingly similar secondary structure shared between these proteins. (D) Regions of sequence conservation in E. coil SecA calculated by Consurf, with the pseudo-helicase motifs in bold revealing their high levels of sequence conservation. (E) A weblogo of E.coli SecA highlighting all the helicase motifs, both nucleotide binding (grey) and translocation (colours) revealing the regions of high sequence conservation. (F) A table displaying the sequences of these identified pseudo-helicase motifs and how they differ from previous identifications by Krishnamurthy et al 2022, Schmidt et al 2001, and Koonin and Gorbalenya 1SS2.

### SecA contains several conserved pseudo-helicase motifs in the proximity of the preprotein and PPXD domain

As SecA is more closely related to SF2 helicases, and as the helicase signature motifs are broadly conserved throughout this branch of the phylogenetic tree, we opted to compare SecA with the Hel308. This Ski2-like helicase was chosen on the basis of its high scoring sequence and structural conservation (Supplementary Data 1 and Supplementary Tables T2 and T3), and its well characterised activity and structure (PDB:2P6R)^33^ (**Figure 2B**). We initially define ‘pseudo-helicase motifs’ as SecA sequences occupying equivalent positions as conventional helicase motifs when comparing the structures of Hel308 to a well-superimposed AlphaFold2^34^ model of *E. coli* SecA (RMSD 3.5Å, **Figures 2B and 2C**). The potential significance of these motifs was further reinforced by their high sequence conservation (**Figure 2D and 2E**). Helicase motifs 1, 2 and 6 (Walker A, B and R-finger respectively) are involved in nucleotide binding and hydrolysis, are highly-conserved across helicases and SecA orthologues, and are essential for SecA activity^30,35^. Therefore, we focus here on the translocation motifs which we have named according to the convention^36^. Pseudo-helicase motifs are generally well-conserved across SecA orthologues, with the sole exception of motif 1b (which presents considerable sequence variation even in conventional SF2 helicases^24^), consistent with their important functional roles. Intriguingly however, and in contrast to the ATP binding and hydrolysis motifs^24^, the translocation motifs (1a, 1b, 1c, 4, 4a and 5) clearly differ in sequence between SecA orthologues and SF2 helicases, presumably reflecting their adaption to a different polymeric client. Some helicase-like translocation motifs have been noted in SecA previously (**Figure 2F)**, and our definitions of the motifs 1a, 4 and 5 are broadly similar^37,38,22^. Importantly though, our approach has significantly extended or defined novel regions of motifs 1b, 1c and 4a compared to the existing literature^38^.

The models of *E. coli* SecA alone (**Figure 2C**) do not represent the active, substrate-engaged, enzyme^6,39,40^. However, a structure of SecA engaged with SecYEG and a short preprotein fragment has been solved for *Bacillus subtilis* (Dong et al; **Figure 3A**), and this was used to generate a homology model with superimposed substrate using SWISS MODEL^41^ (**Figure 3B**). In this model the interactions of the helicase motifs and preprotein are evident^6^. Preprotein sits above NBD1, passing over motif 1c, and close to the beta strands that lead to the PPXD domain. The N-core motifs in helicases are usually attributed to substrate binding and specificity: motif 1a is often involved in translocation, motif 1b in specificity and unwinding and motif 1c also in unwinding^24^. In SecA, motif 1a is furthest from the substrate and includes highly conserved N132 and D133 at the tip of the helix pointing towards the substrate cleft (**Figure 3C**). In motif 1b, there is a weakly conserved proline (P159) that sits at the apex of the loop (**Figure 3C**) similar to the structure of other DEAH/RHA ‘hook-turn’ 1b motifs^32^. Finally, motif 1c contains two key regions including asparagines (N179 and N180; **Figure 3C**) which protrude towards the substrate, and a more C-terminal phenylalanine (F184), which contacts the substrate^6^ (**Figure 3C**).

**Figure 3:**
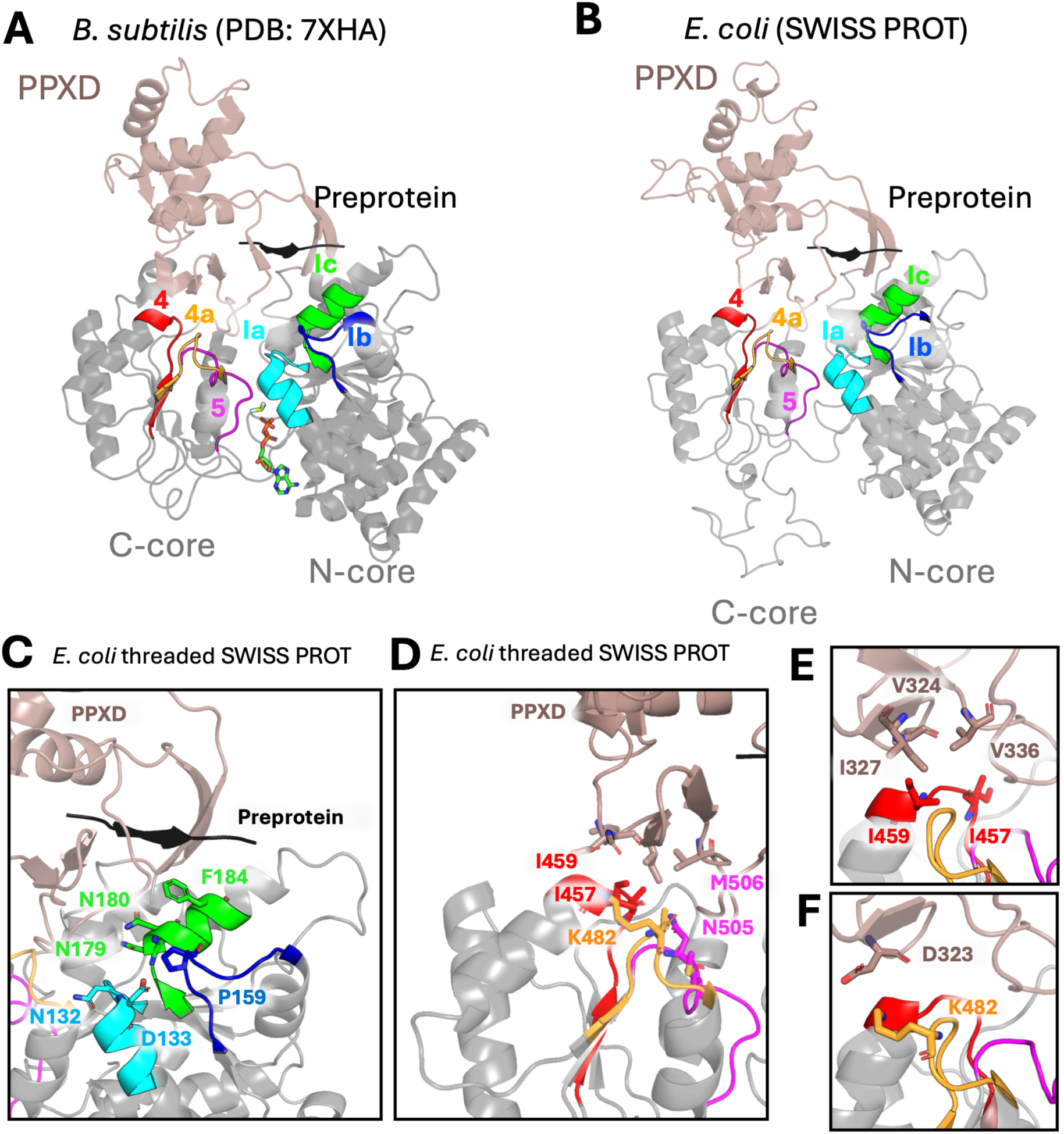
Pseudo-helicase motifs within SecA. (A) The B. subtilis structure (PDB:7XHA)^C^ with the resolved location of preprotein. The pseudo-helicase motifs are shown in colours, and the PPXD domain is shown in light brown, clamped down onto NBD2. (B) A model of the E. coli sequence made using SWISS MODEL, revealing a predicted position of the PPXD domain, shown in light brown, above the NBD2 motifs. The location of the resolved preprotein from the B. subtilis structure is superimposed into this model and shown in black. (C) A zoom into the NBD1 pseudo-helicase motifs of the E. coli SWISS MODEL depicting the three pseudo-helicase motifs, 1a shown in cyan, 1b shown in blue and 1c shown in green, the phenylalanine in motif 1c has been identified for contacting the substrate in B. subtilis^C,40^. The key conserved residues of each motif are depicted by sticks. (D) A zoom into the NBD2 pseudo-helicase motifs of the E. coli SWISS MODEL, with the pseudo-helicase motifs in bold, motif 4 is shown in red, motif 4a in orange, and motif 5 in magenta. The key conserved residues of each motif are depicted as sticks. (E) The interface between pseudo-helicase motif 4, shown in red, and the PPXD domain, shown in light brown, is formed of hydrophobic interactions between both I457 and I45S with hydrophobic residues on the PPXD domain, collectively forming a hydrophobic interface that is highly conserved on both domains. The residues involved in this interaction are depicted with sticks. (F) The interface identified between pseudo-helicase motif 4a, shown in orange, and the PPXD domain, shown in light brown, is made up of a salt bridge. The residues involved in this interaction are depicted with sticks.

A key distinction between SecA and SF2 helicases is the PPXD, which is unique to SecA (Supplementary Tables T4 and T5, Supplementary Data 2). The PPXD is a dynamic preprotein binding region that undergoes conformational changes upon the addition of ATP^9,42^. Interestingly, in the substrate-bound structures of SecA (**Figure 3B**), the PPXD sits immediately above NBD2 in very close proximity to pseudo-helicase motifs 4, 4a and 5 and appears to clamp down against them on the addition of substrate^6,40,43^. Motif 4 comprises a loop into a short turn of helix with two highly conserved hydrophobic residues (I457 and I459) that protrude into the space between PPXD and NBD2 (**Figure 3D**). Motif 4a is primarily made up of a loop that contains a highly conserved charged residue (K482), which reaches into the PPXD:NBD2 interface (**Figure 3D**). Finally, pseudo-helicase motif 5 sits at the interface between NBD2 and NBD1, and contains the highly conserved residues N505 and M506, near the nucleotide binding cavity (**Figure 3D**).

To investigate the interactions across this PPXD-NBD2 interface (**Figure 3D**), ePISA^44^ and EV-couplings^45^ were used to find residues which interact or display strong co-evolution signatures (Supplementary Data 3 and 4). This identified high confidence interactions and co-evolution in a conserved hydrophobic patch involving: I457 and I459 (motif 4) respectively binding to V324 and I327/V336 (PPXD) (**Figure 3E**). This was in addition to a highly co-conserved salt bridge between K482 (motif 4a) and D323 (PPXD) (**Figure 3F**). In SF2 helicases, motifs 4 and 4a directly grip and release DNA substrates in response to the ATP hydrolysis cycle to promote translocation^24^. By analogy, our observation here implies that the PPXD domain (and *indirectly* preprotein) may undergo controlled remodelling in response to ATP binding and hydrolysis mediated through motifs 4 and 4a.

### The NBD1 pseudo-helicase motifs regulate ATPase activity and its activation by preprotein

To investigate the role of the NBD1 pseudo-helicase motifs in SecA (1a, 1b and 1c), amino acid substitutions were designed for each of the highly conserved residues identified above. Purification of all variants proceeded just as native SecA, and appeared to be fully folded according to CD spectra (Supplementary Figure S1).

We next performed SecA-SecYEG ATPase assays in the absence and presence of preprotein using a coupled spectrophotometric assay (**Figure 4**). Proteoliposomes were provided in excess to saturate the added SecA (see Methods for details). As expected^13,46^, the native SecA-SecYEG displayed very low ATPase activity (∼ 0.3 s^-1^) that was stimulated (∼ 20-fold) upon the binding of preprotein substrate. This stimulation is thought to result from allosteric destabilisation of the SecA:ADP complex^38,47^. The N132A/D133A variant protein (motif 1a) showed similar SecY-stimulated ATPase activity but dramatically reduced stimulation by preprotein (**Figure 4A/E**). While the P159A variant (motif 1b) displayed a ∼ 3 fold increase in SecY-stimulated ATPase activity, but was also poorly stimulated by preprotein (**Figure 4B/E**). The N179A/N180A variant (motif 1c) did not change the SecY-stimulated rate but dramatically reduced preprotein stimulation (**Figure4C/E**). Finally, the F184A variant (motif 1c) significantly reduced SecY-stimulated ATPase and essentially eliminated stimulation by preprotein (**Figure 4D/E**). Overall, the most striking result was that all variants in motifs in NBD1 dramatically reduce ATPase stimulation by preprotein, indicating a critical role for this domain in sensing or communicating the presence of substrate to the ATPase site, and its subsequent activation. These data also suggest regulatory roles for P159 (motif 1b) and F184 (motif 1c) in the SecY-stimulated ATPase rate in the absence of preprotein.

**Figure 4:**
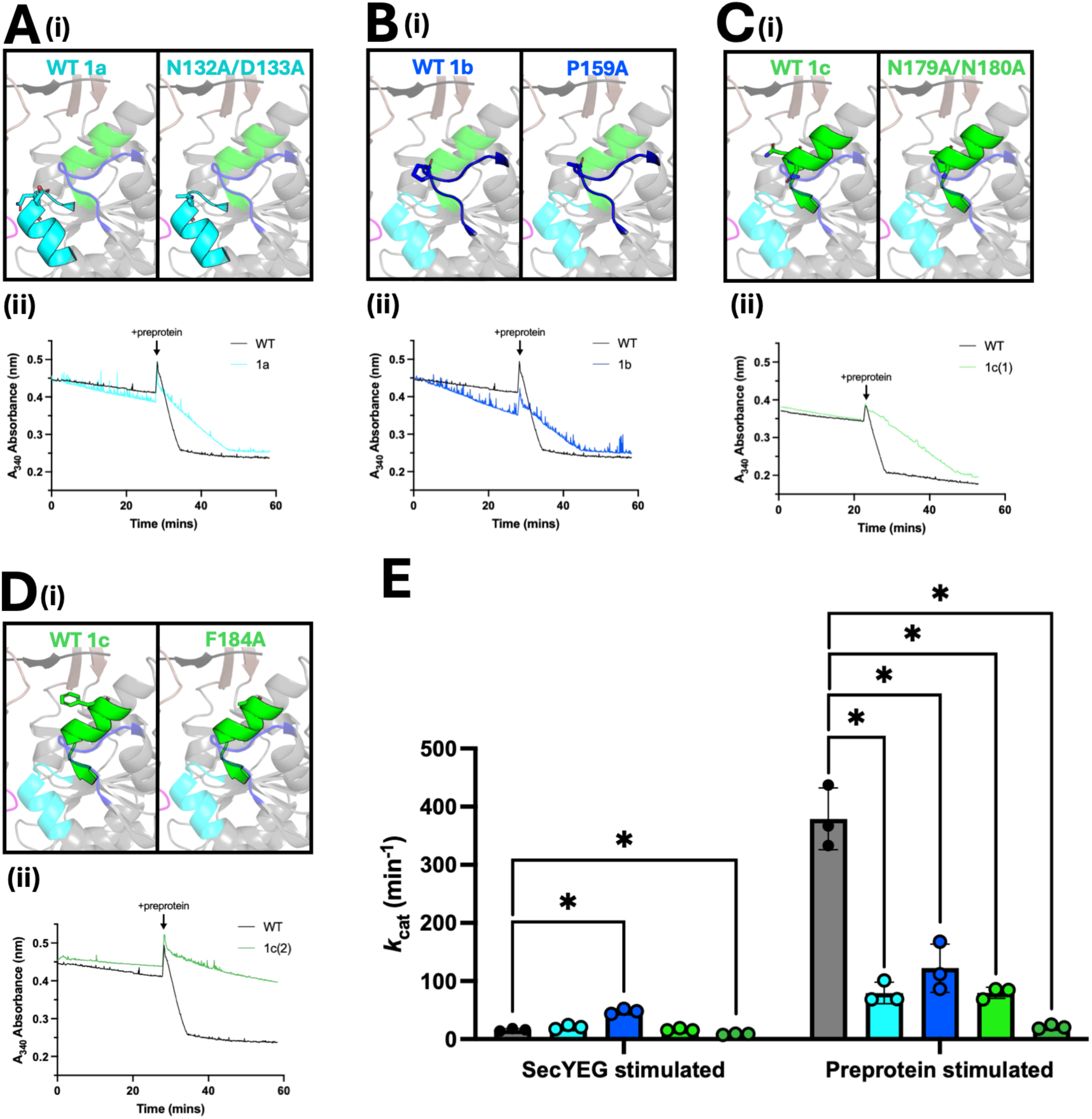
NBD1 pseudo-helicase motifs impact ATPase activation in SecA. (A) The amino acid substitutions made in motif 1a are depicted with sticks, alongside an example ATPase assay trace. (B) Motif 1b; (C) The N-terminal amino acid substitution in motif 1c; (D) The C-terminal amino acid substitution in motif 1c. (E) Comparison of the ATPase rates (kcat) for each amino acid variant when SecYEG-stimulated, or preprotein-stimulated, with significantly different rates indicated with a “*”.

### Pseudo-helicase motif 1c is critical for protein transport

We next investigated protein translocation by SecA variants using an assay based on a split NanoLuc luciferase^48,49^ (see Methods). In these assays the large fragment of a luciferase (11S) is encapsulated in the proteoliposome interior and the complementary small fragment (pep86) is fused to the C-terminus of the precursor. Once pep86 contacts 11S a luminescent signal is generated as a measure of preprotein transport. As translocation proceeds N→C the pep86 enters last, so that a lag is generated that reflects the time taken for the preceding preprotein to be transported. Thus, the lag provides an accurate measure of the transport activity: the time taken to initiate transport and progress the pep86 to the proteoliposome interior for association with 11S . Whilst the amplitude is proportional to the maximal amounts (yield) of transported precursor.

Surprisingly, amino acid substitutions in motifs 1a and 1b (N132A/D133A and P159A) and the N-terminal region of motif 1c (N179A/N180A) did not reduce preprotein translocation compared to native SecA (**Figures 5A-C**). By comparison, the preprotein-stimulated ATPase activity was substantially reduced in all these variants (**Figure 4**). Evidently then, similar rates of translocation can be achieved with lower rates of ATP turnover. This varying sensitivity of ATP hydrolysis and transport activities suggest that motif variation is affecting the coupling process. In contrast to this, a variant of the C-terminal region of motif 1c (F184A), reduces both ATPase and preprotein translocation rates (**Figure 4 and 5D**), consistent with previous data showing F184 directly contacting the translocating substrate^6,40^. Variations of the transport yield relative to ATP is difficult to explain; clearly, the involvement of the helicase motifs in ATP driven transport is complex and requires further analysis.

**Figure 5:**
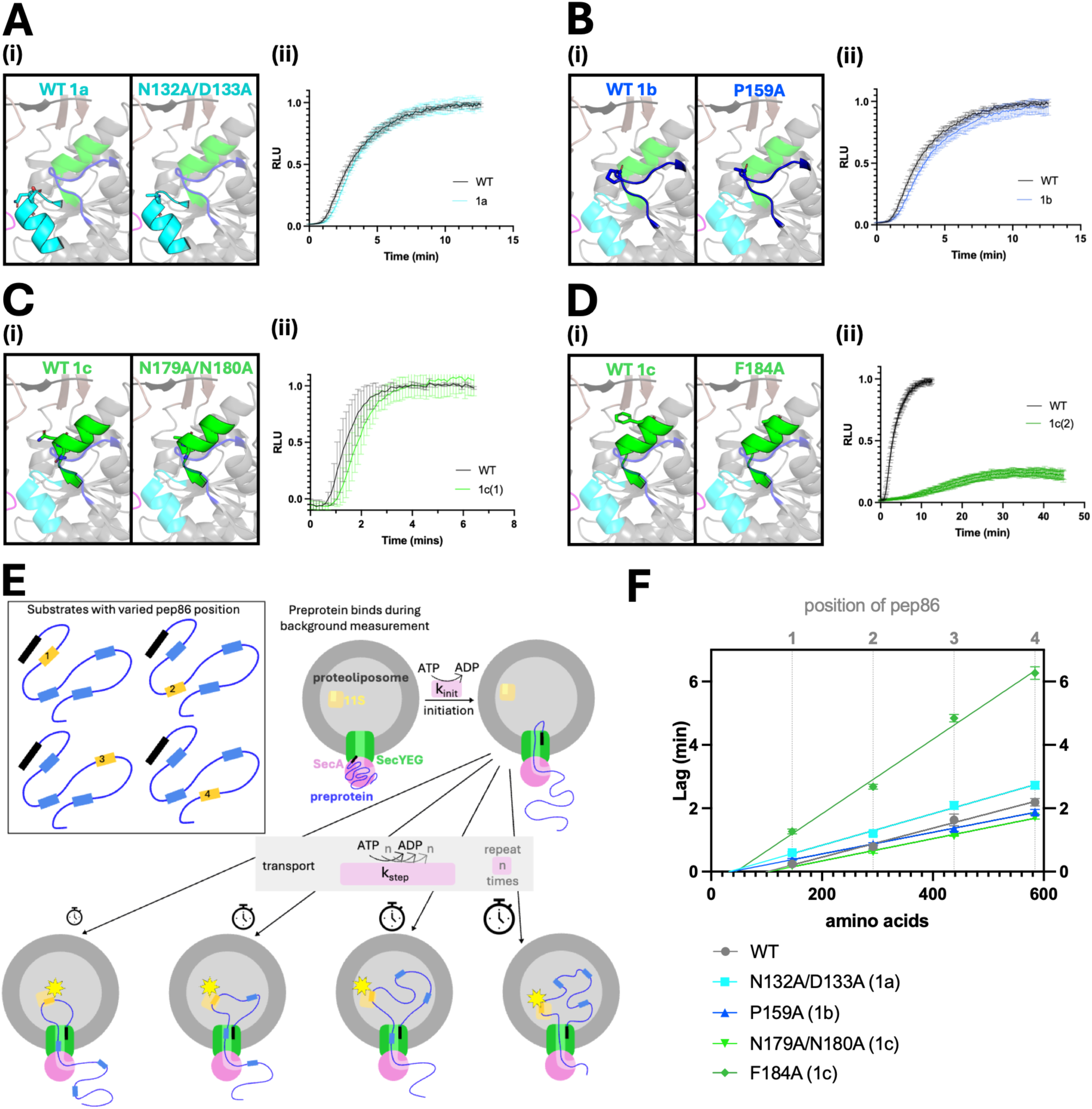
NBD1 motif 1c is important for protein translocation. (A) The amino acid substitutions made in motif 1a are depicted with sticks, alongside the NanoLuc data. (B) Motif 1b; (C) The N-terminal substitution of pseudo-helicase motif 1c; (D) The C-terminal amino acid substitution in motif 1c; (E) Lag experimental design, the small fragment (pep8C) is positioned at increasing distances from the N-terminus of the preprotein, with longer lengths requiring more time to be transported into the lumen of the proteoliposome to encounter 11S and produce a luminescent signal. (F) The lag times for the transport of different lengths of preprotein via the different protein varients. Most of the protein varients (1a – cyan, 1b – blue, and 1c(1) – bright green), showed very similar trends to native SecA (grey), with a linear increase in lag time corresponding with increasing preprotein length, with the variant in the C-terminal region of motif 1c (1c(2) – dark green), showing the most different trend. Linear regression lines were generated for each data set and extrapolated to the x-intercept.

To dig further, the respective SecA variants were tested with a series of substrates in which the pep86 fragment is placed at increasing distance from the signal sequence (**Figure 5E**) in order to quantify the translocation kinetics^11^. The time it takes for the substrate to be translocated to the pep86 fragment is given by the lag time, which can be accurately measured for increasing stretches of polypeptides (**Figure 5E**). In a plot of amino acid length versus lag time (**Figure 5F**), the gradient determines the rate of translocation (∼ 220 amino acids/min for native SecA), and the X-axis intercept relates to the initiation of transport^16^.

In this assay the preprotein is engaged with SecA and the translocon prior to the addition of ATP, which then stimulates initiation and the flipping of the signal sequence into the lateral gate of SecY, bringing with it a region of mature domain^11^, prior to translocation. The translocation rate for the N132A/D133A variant (motif 1a) is not significantly affected (∼200 amino acids min), but has a less effective initiation (smaller intercept with the x-axis). Similarly, the P159A (motif 1b) and N179A/N180A (motif 1c) variants bring about a slightly faster rate of transport, respectively ∼ 290 amino acids/ min^-1^ (∼ 1.3-fold faster) and 280 amino acids/min (∼ 1.3-fold faster), with the former exhibiting a less effective initiation (smaller intercept with the x-axis). In stark contrast, variant F184A (motif 1c) displays a very different profile: a much slower translocation rate (85 amino acids/ min, ∼ 0.4-fold) and less effective initiation.

The above results identify distinct effects for NBD1 variants, particularly the C-terminal variant of motif 1c (F184A), identifying a crucial role for this region in the coupling process, potentially through its known substrate association (**Figure 3C**)^6,40^.

### The NBD2 pseudo-helicase motifs are essential for productive engagement of preprotein

Next, we investigated the role of PPXD interaction motifs in NBD2 for the activation of ATPase activity. Again, all protein variants were purified in the same manner as native SecA, and displayed normal CD spectra indicating correct folding (Supplementary Figure S1). The ATPase activity of I457Ǫ/I459Ǫ variant (motif 4) was stimulated by SecYEG (in proteoliposomes) alone, similar to native SecA, but not further enhanced by preprotein as it should have been (**Figure 6A, 6E**). Similar results were obtained for variants A481S/K482S and A481Ǫ/K482A (motif 4a) (**Figure 6B, 6C, 6E**). In contrast though, ATP turnover of the N505A/M506A (motif 5) variant was excessively activated by SecYEG alone, and then further stimulated by preprotein, albeit to a lesser extent than native SecA (**Figure 6D, 6E**).

**Figure 6:**
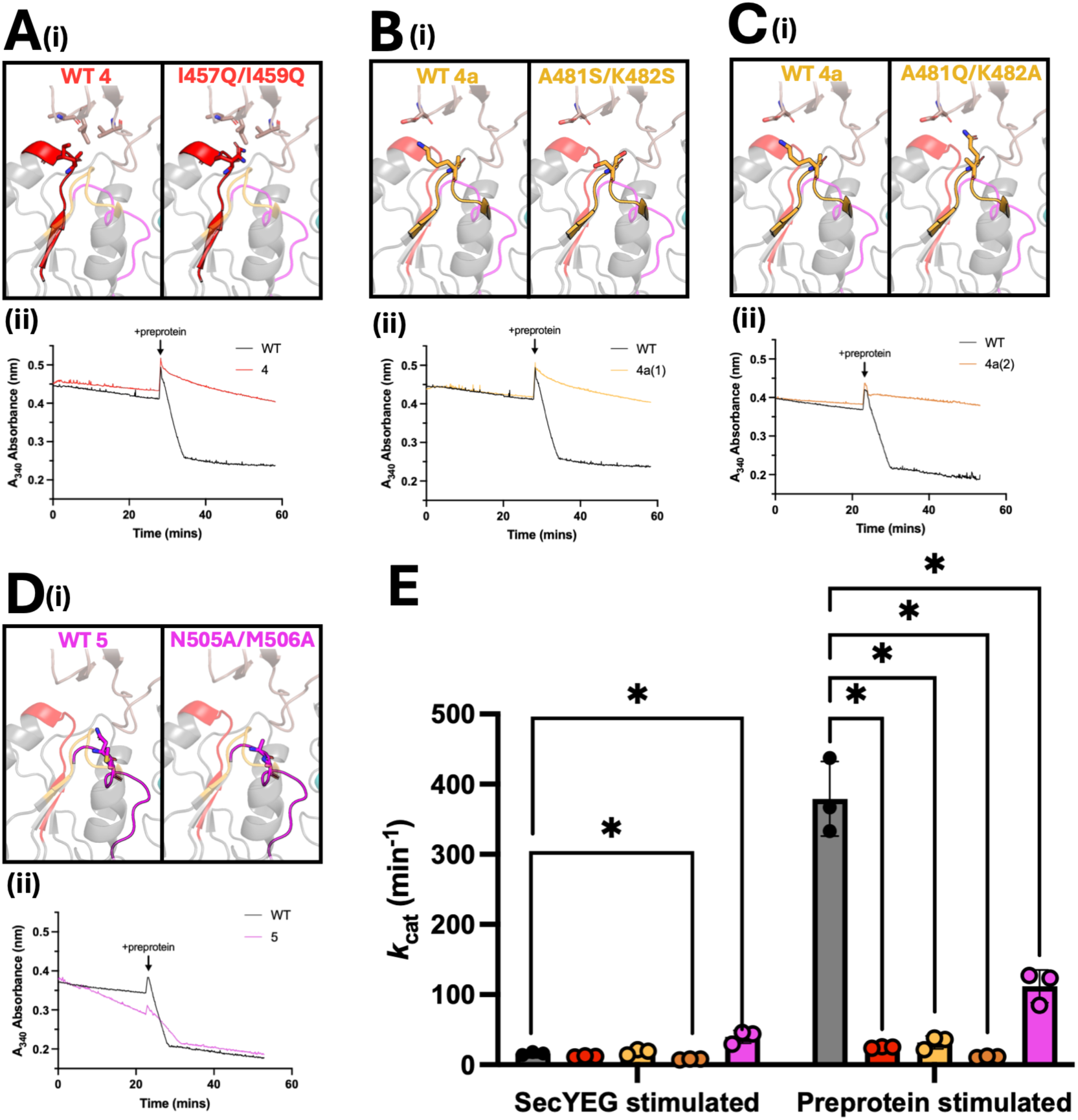
NBD2 pseudo-helicase motifs are essential for adequate ATPase stimulation on the addition of preprotein. (A) Closeup of motif 4, with substituted residues (native left and substituted right) depicted as sticks (i), with an example ATPase trace (ii). (B-D) As in panel A, but for (B) The first substitution in motif 4a(1); (C) The second substitution in motif 4a(2); and (D) motif 5.(E) Comparison of the ATPase rates (kcat) for each amino acid variant when SecYEG-stimulated, or preprotein-stimulated. The protein variants that resulted in significantly different rates are indicated with “*”.

Notably, the NBD2 variants (with the exception the motif 5 version, which potentially directly influences the NBS (**Figure 3A**), all had lower preprotein stimulation than the NBD1 variants (compare **Figure 6E** **with 4E**). Thus, it appears that motifs 4, 4a and 5 are crucial for the allosteric communication which connects substrate binding and activation of ATPase activity. This also highlights the importance of the interaction between the NBD2 and PPXD domain *via* K482 (**Figure 3D**) in SecA SecY-stimulated ATPase activity.

### The NBD2 pseudo-helicase motifs at the PPXD interface are essential for protein transport

We next tested all NBD2 variants for their ability to transport protein using the NanoLuc assay. The I457Ǫ/I459Ǫ (motif 4) and A481S/K482S (motif 4a) variants both showed an almost complete loss of transport activity (**Figures 7A, 7B**). The A481Ǫ/K482A variant (motif 4a) showed a reduction in transport activity (longer lag time) and approximately half the signal amplitude (transport yield) seen in native SecA (**Figure 7C**). Finally, the N505A/M506A variant (motif 5) exhibited slower transport (longer lag), but a transport yield similar to the native enzyme (**Figure 7D**).

**Figure 7:**
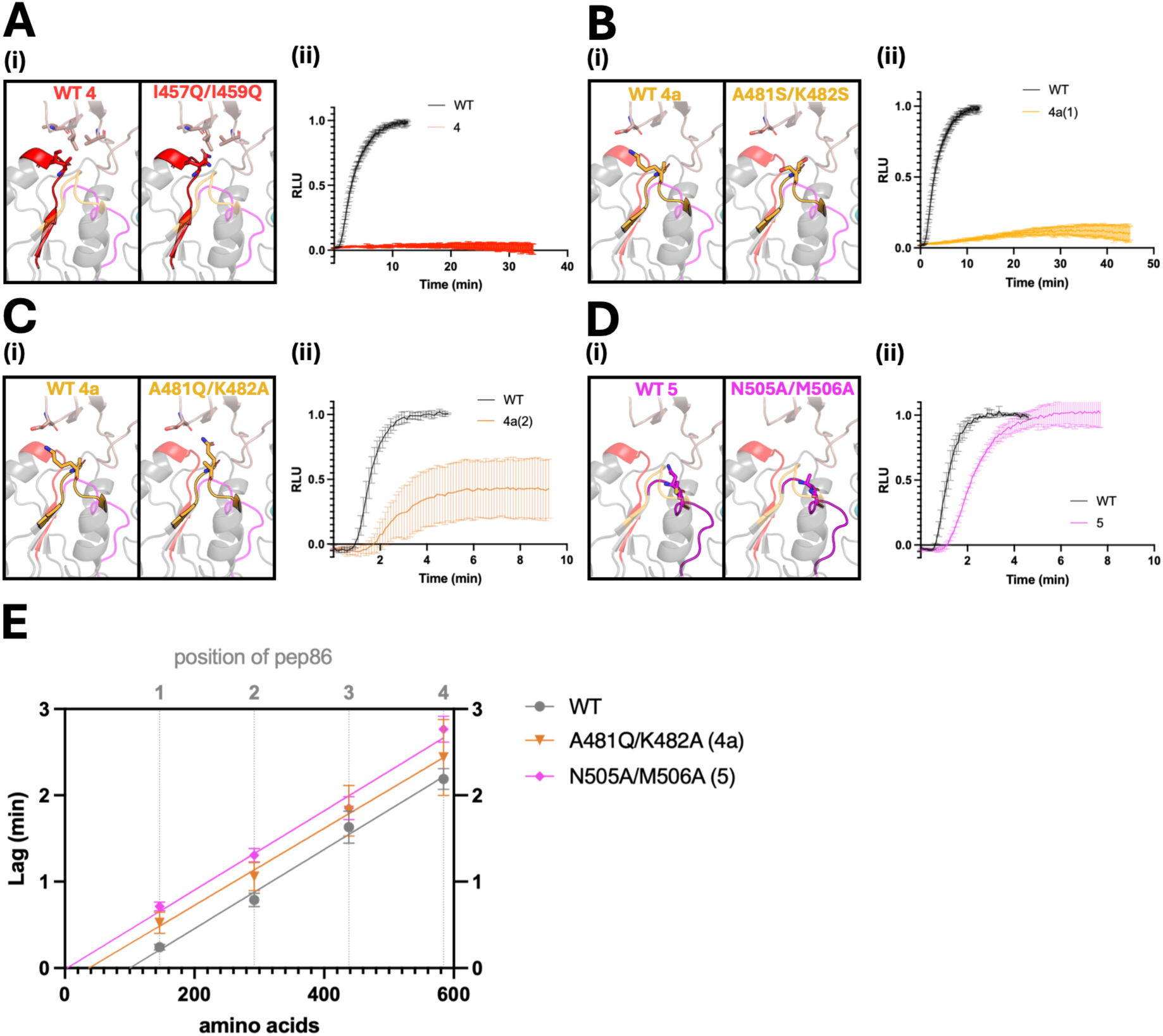
The NBD2 pseudo-helicase motifs at the PPXD interface are essential for preprotein transport. (A) The amino acid substitution made in motif 4 is depicted with sticks, alongside the NanoLuc data, shown in red and depicting an inability for this protein variant to transport; (B) the amino acid substitution with serines in motif 4a;.(C) the amino acid substitutions to glutamine and alanine made in motif 4a; (D) motif 5. (E) The lag times for the transport of different lengths of preprotein via the different protein varients, excluding the protein varients in 4 and 4a(1) that fail to transport adequately to calculate accurate lag times. The protein variants 4a(2) and 5 show a linear trend with the same gradient as native SecA. Linear regression lines were generated for each data set and extrapolated to the intercept. Both 4a(2) and 5 intercept closer to the y-axis, compared to native.

For the latter two, which retained transport activity, additional experiments were performed with pep86 fragments at different positions relative to signal sequence (**Figure 7E**). For both A481Ǫ/K482A and N505A/M506A variants there is a simple linear relationship between lag time and preprotein length with the same gradient as native SecA, suggesting translocation at the native rate (∼ 220 amino acids/ min). However, both protein varients show a consistently longer lag for each pep86 position, and an X-intercept shifted closer to 0, likely due to slower initiation.

## Discussion

In this work we investigated the potential functional similarities between protein translocase SecA and SF2 nucleic acid helicases. Phylogenetic and structural analysis of SecA indicates that it is most similar to SF2 helicases. On this basis, we defined pseudo-helicase motifs in SecA, analysed their substrate and intra-domain interactions in existing structures/models, and experimentally tested their importance for ATPase activity and coupled protein translocation using a biochemical approach involving the reconstitution and monitoring of ATP-driven protein transport. The Walker A, Walker B and arginine finger motifs are widely shared across vast numbers of ASCE P-loop proteins and their roles in ATP binding and hydrolysis are well-characterised^23,25^. Therefore, our work focused instead on those conserved SecA motifs that are structurally equivalent to the nucleic acid binding and translocation motifs found in the helicases.

Most NBD1 motif mutations affect coupling between preprotein binding and SecA ATPase activity, albeit in subtly different ways. However, under the conditions of our assay, they transport protein at nearly native rates regardless; though this may not be the case for more challenging preprotein clients. These motifs localise to the upper surface of NBD1, and in some cases make direct contact with the substrate polypeptide (**Figure 8**). Our results are reminiscent of similar findings in SF1 and SF2 helicases where ATPase stimulation by nucleic acids is regulated by conserved N-core motifs^50–53^. Disrupting conserved features in motif 1a produced lower stimulated ATPase rates and apparently less effective initiation, but native levels of processive protein transport. Additionally, perturbations to motif 1b resulted in a significantly higher SecY-stimulated rate of ATPase activity in the absence of preprotein. The substitution of P159A at the apex of a loop protruding from NBD1 towards the substrate binding site, both alleviates suppression of ATPase activity in the absence of substrate and interferes with ATPase activation in its presence. This amino acid substitution may have disrupted a conformation of the loop that could normally lock SecA in the ADP state in the absence of preprotein, resulting in a higher SecY-stimulated rate, while also preventing substrate-induced allostery that unlocks this state to promote the higher translocation-coupled ATPase activity. Interestingly, conformational changes in the 1b loop are associated with SecYEG binding and ATPase stimulation^38^ (**Figure 8**). The SecYEG and preprotein stimulated ATPase rates are also sensitive to disruptions to motif 1c, consistent with the idea that this motif allosterically regulates the conformation of the NBS(**Figure 8**). For example, the mutation F184A in this motif reduced SecYEG-stimulated ATPase rate and prevented the further activation by preprotein. Given that this residue interacts directly with substrate^6,40^, our work implies that motif 1c is also part of an allosteric network allowing the ATPase to respond to substrate and *vice versa* (**Figure 8**).

**Figure 8:**
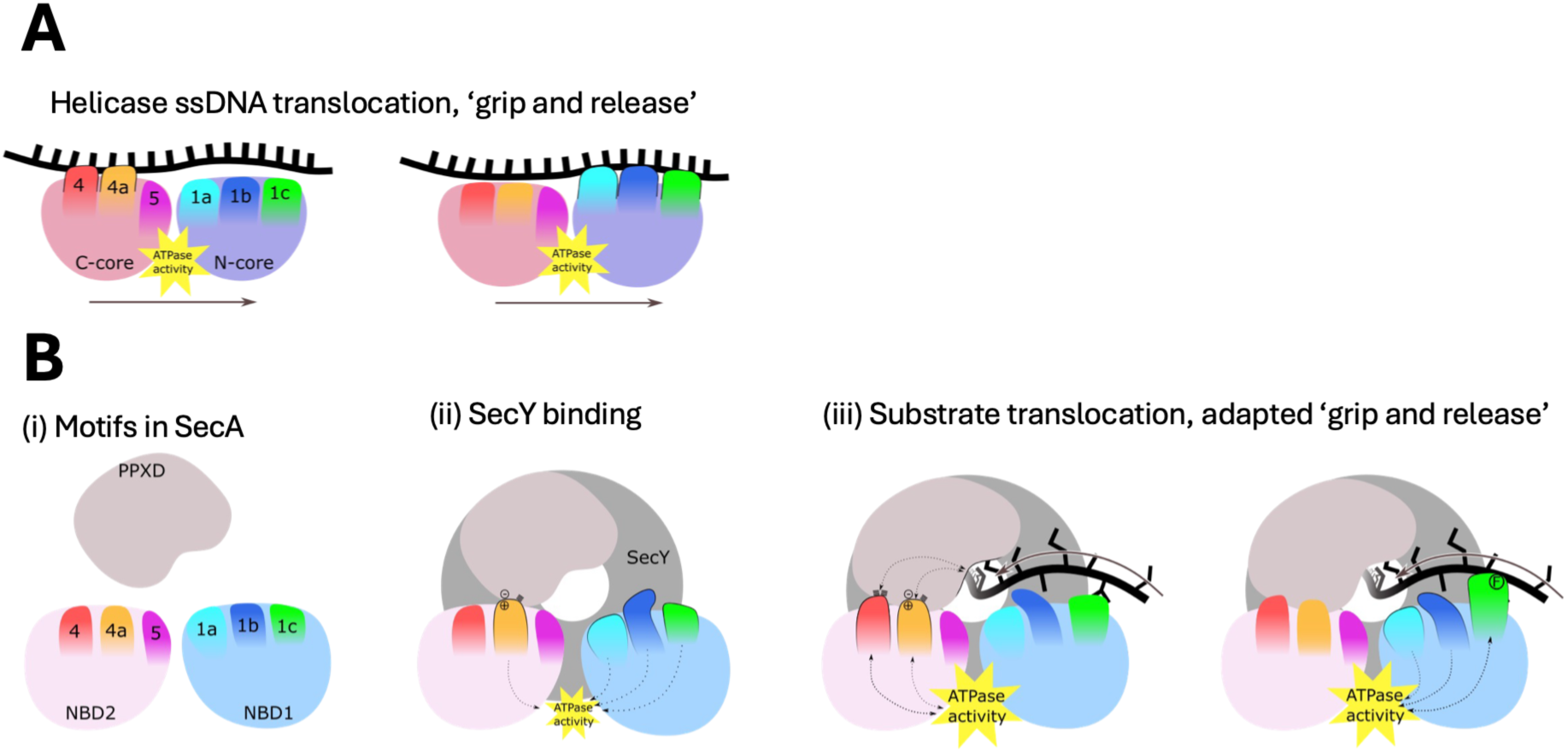
Summary of the responses seen in the pseudo-helicase motifs in SecA. (A) A representation of a SF2 helicase translocating ssDNA via the ‘grip and release mechanism’. (B)(i) The pseudo-helicase motifs in SecA, (ii) Upon SecY binding, motif 1b undergoes a conformational change, and the NBD1 motifs stimulate a low level of SecY-stimulated ATPase activity, NBD2 motif 4a may sense the movement of the clamp upon SecY binding (iii) Substrate binding drives the PPXD domain to clamp down on top of NBD2, where this is sensed by the NBD2 pseudo-helicase motifs, that also coordinate between the substrate binding site on PPXD and the ATPase activity. We propose that this occurs through allosteric networks via the conserved interactions between PPXD and NBD2 through the hydrophobic patch interaction of motif 4 and the salt bridge and H-bond network in motif 4a. This co-ordinates the high rate of ATPase activity during translocation with the PPXD substrate contacts and potential remodelling. The NBD1 motifs also respond to substrate with the phenylalanine in motif 1c making direct substrate contacts, and a proposed conformational change in motif 1b to further stimulate ATPase activity for translocation. The NBD1 motifs communicate the presence of substrate to stimulate high levels of ATPase activity associated with translocation. In summary we postulate that these pseudo-helicase motifs could facilitate preprotein translocation via a similar ‘grip and release’ mechanism, but rather using the PPXD as an adaptor for contacting preprotein.

Additionally, variants with substitutions in motif 1b and the N-terminal region of 1c retained almost native-like transport suggesting these regions play no direct role in preprotein transport, and revealing a lack of tight coupling between ATPase and transport activity. In fact, these variant proteins are apparently more efficient than native SecA in this experimental context. Although superficially surprising, our results are consistent with observations that there is loose coupling between the ATPase activity and translocation through the Sec system^11,17^, with additional support coming from the rapid opening of the SecY channel that occurs much faster than ATP hydrolysis^54^. This is entirely consistent with the proposed Brownian ratchet mechanism of ATP driven protein translocation^4,17,55^. In the context of which, it will be interesting to see if these SecA variants exhibit more nuanced defects during the transport of polypeptide *in vivo*. Or, if they struggle to transport precursors which place higher demands on ATP-driven movement versus diffusion, such as those containing highly positive, hydrophobic or bulky residues.

In contrast, the F184A amino acid substitution within motif 1c severely reduced transport. This indicates a more direct mechanistic role in moving polypeptide (**Figure 8**,consistent with the role of motif 1c in making substrate interactions), and/or its reduced substrate-stimulated ATPase activity is now too low to achieve native-like transport. The varying ATP efficiency seen in the protein varients seems incompatible with a power-stroke mechanism.

The pseudo-helicase variants in NBD2 (motifs 4, 4a and 5) have little effect on SecYEG-stimulated ATPase activity, but those at the interface with the PPXD dramatically reduce the preprotein stimulated ATPase and, with the exception of motif 5, are devoid of preprotein transport. This is not unexpected, as this interface connects the helicase-core of SecA to the PPXD substrate binding domain (**Figure 8**). Thus, the results suggest a crucial role for the helicase motifs at the NBD2-PPXD interface in the propagation of ATP-dependent transport, potentially through effects on substrate binding or remodelling. This idea is supported by existing work suggesting that the region just beyond the 4a motif exhibits preprotein-dependent dynamics: binding of preprotein drives the PPXD to clamp onto NBD2, with increased dynamics in the motif regions, enhancing the ATPase activity^47^. This same study revealed an H-bonding network between regions of NBD2 and the PPXD, including residue H484 in motif 4a. Together, the observations made in our study, and elsewhere^47,56^, suggest a model where the PPXD domain insertion into the helicase core adapts the normal substrate ‘grip and release’ mechanism for ssDNA translocation, executed by the helicase motifs on the C-core, for the unique function of SecA in translocating protein substrates through a pore (**Figure 8**). Indeed, inserts within the helicase domains are used broadly to adapt force generation for a particular substrate or function. For example, the Swi/Snf family helicases use a unique domain to transduce ATPase to translocation and rotational force required for chromatin remodelling activities^57^. Similarly, XPD helicases exploit a unique domain to enable the recognition of different DNA lesions to enact optimal repair strategies^58–60^. The lower SecYEG-stimulated ATPase activity in the A481Ǫ/K482A amino acid substitution in motif 4a could reflect its involvement in sensing the closure of the clamp between PPXD-NBD2 (**Figure 8**), required for stimulation of ATP-turnover, preprotein capture and coupled translocation^9,43,61^. Overall, the results show that pseudo-helicase motifs in the NBD1 domain regulate SecYEG-stimulated ATPase activity and mediate substrate sensing for ATPase activation whereas the NBD2 motifs appear to co-ordinate communication with the PPXD domain for coupling of ATPase to peptide transport.

Following our classification of SecA as a SF2-like ‘helicase’ we went on to explore if there might be other adapted helicases which convert energy from ATP hydrolysis into mechanical work on non-conventional substrates (Materials and methods). However, SecA and the closely related SecA2 proteins^62^ appear to be unique in their use of an adapted SF2 helicase core for protein transport, with both containing the PPXD domain insert in their helicase core (Supplementary Table T6). It is noteworthy, however, that SF2 helicases and helicase-like proteins themselves display extraordinary diversity in terms of their substrate specificity and function, albeit usually directed towards nucleic acids rather than polypeptides. These enzymes target both RNA and DNA, can exert their activity on either single- or double-stranded nucleic acids, can move unidirectionally with either 3’-5’ or 5’-3’ polarity, and in some cases can act as chemo-mechanical switches rather than processive motors^24,63^. Therefore, although SecA has been considered a complete outlier, one might instead view its behaviour as part of the continuum of diversity observed for all SF2 helicases. The most basic function shared by all these molecular machines is their ability to transduce energy derived from ATP turnover into some form of mechanical manipulation of the substrate. It is this underlying function that is associated with the core domains, where Walker A, B and the arginine finger (helicase motifs 1, 2 and 6) surround the ATP molecule promoting its hydrolysis when the bipartite core adopts a compact closed form. In turn, hydrolysis drives core opening and other localised conformational changes to manipulate the substrate. Studies of conventional helicases revealed that it is other motifs (including 1a, 1b, 1c, 4, 4a and 5) that play a key role in the allosteric communication between the ATPase cycle and the substrate remodelling, with many of these conserved amino acids directly involved in substrate contacts. It is no surprise then that, whereas motifs 1, 2 and 6 are strongly conserved across all the diverse members of SF2, it is these substrate-specific motifs in which we observe the greatest diversity. This is illustrated throughout several helicase superfamilies containing proteins that interact with different substrates and in different ways, whilst sharing mechanistic features that group them as a family^27^. It is likely that SecA has arisen from an ancient helicase-like common ancestor that contained the core helicase domains, from which specialised substrate interaction motifs evolved alongside the insertion of the PPXD. More broadly, this suggests that diverse ATP-driven molecular machines may share conserved catalytic engines, repurposed through evolution. What distinguishes them may instead be their substrate-interacting elements, which dictate the nature and direction of their mechanical work.

## Methods

### Phylogenetic analysis

Representative sequences of each helicase subfamily within SF1 and SF2 were extracted from a previously-compiled dataset of 700 representative bacterial and archaeal proteomes^64^. For each subfamily, a profile hidden Markov model representing the subfamily was built with hmmbuild (HMMER v3.4^65–67^ program) from an alignment of the subfamily members identified from a previous phylogenetic study^24^. These subfamily sequence profiles were then used to search (hmmsearch) the large and diverse dataset to generate the SF1 and SF2 representative sequences dataset. From the ranked significant hits of each search, the top 20 bacterial and top 20 archaeal sequences, each from a different organism, were selected. Sequences identified by more than one subfamily profile were incorporated into the data set once and labelled with all relevant subfamilies. The final generated dataset contains 565 sequences. Sequences were aligned with MAFFT L-INS-i^68,69^ and maximum likelihood phylogenies were inferred from the MAFFT alignment (565 sequences, 9,217 aligned positions) with IǪ-TREE2^29^ 2.4.0. IǪ-TREE was used along with the ultrafast bootstrap^70^ to generate an evolutionary tree. The best fit substitution model, Ǫ.pfam+I+R10, was selected using ModelFinder. Branch support was assessed with 1,000 ultrafast bootstrap replicates and 1,000 SH-like approximate likelihood ratio test replicates.

### Sequence and structural analysis

The solved structure of *B. subtilis* SecA (PDB: 7xha) was used for structural searches in both the Foldseek webserver^71^ with the standard search parameters, against all databases, and the DALI webserver^72^ using the PDB search, identifying many high confidence helicase hits. After validating the structural helicase comparison through these searches a well-defined helicase Hel308 was used for structural comparisons. We generated a *de novo* resting structure of E. coli SecA using AlphaFold2^34^. To generate an active conformation we used SWISS-MODEL^41,73–77^, with a solved structure (PDB code 7xha) as a template. Comparisons were then made in PyMOL. For the sequence conservation studies, the ConSurf webserver^78–83^ was used to generate a structure with each residue scored by degree of evolutionary conservation, and WebLogo version 3.7.12^84^ was used to visualise the multi-sequence alignment generated by the ConSurf webserver. EVCouplings^45^ was used to investigate the co-evolution of sites in SecA, sites with probability scores over 0.9 were defined as confident. The PDBePISA^44^ webserver (EMBL-EBI), interface analysis was also used to find interacting residues. To do this the PPXD and NBD2 domains were extracted from PDB:7xha^6^ in PyMOL and saved as separate chains so PISA could analyse their interface, (PPXD as structure 1 and NBD2 as structure 2). For sequence analysis a profile hidden Markov model for SecA was generated using hmmbuild (HMMER v3.4) from the top 10 representative SecA bacterial sequences, from different organisms in the 700 proteomes dataset, followed by hmmsearch to search for other proteins that may have similarities in their sequences, revealing helicase hits only to the NBDs (Supplementary Tables T2 and T3).

### Protein purification

All reagents reported previously for the assays were produced as described previously^49^, SecYEG^85^, pSpyXLX^55^, pSpyL/D series^11^, and proteoliposomes^86^. The SecA protein variants were generated by site-directed mutagenesis using the ǪuikChange method (Agilent), primers from Eurofins Genomics, and prepared as for wild type^86^. SecA was prepared using our standard protocol. The SecA variants were purified in the same way but following a 3hr expression, and tested for misfolding using Circular Dichroism.

### Circular Dichroism spectroscopy

To measure the circular dichroism (CD) spectra a JASCO J-815 CD polarimeter was used. Sample were loaded at 2 µM in 20 mM Tris/HCl pH7.5, 50 mM KCl, 2 mM MgCl2 (buffer used to provide a baseline for the protein spectra) into a 1mm pathlength quartz cell. Full CD spectra were measured between 190-250 nm at regular temperature intervals during a 5-95 °C melt, acquiring eight spectra at each temperature and averaging the results. The raw circular dichroism data were converted into mean residual ellipticity (MRE) using the following equation (where n is the number of peptide bonds in a protein sample):

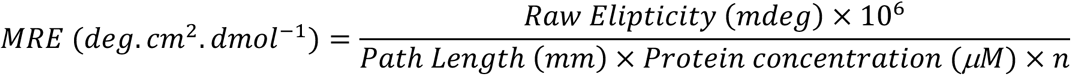

### ATPase activity assays

Steady-state ATPase assays, with the consumption of ATP coupled to the oxidation of NADH, were carried out as in previous studies^86^. This oxidation is coupled using an enzyme system containing pyruvate kinase and lactate dehydrogenase (PK/LDH) (Merck) with excess phosphoenolpyruvate (PEP). The absorption of NADH at 340 nm was measured in a BioTek Synergy Neo2 plate reader.

For each set of measurements, a master mix was made up in TKM to reach a final volume of 84 µl with 1 µl PK/LDH, 230 nM SecYEG in proteoliposomes, and other components added to give final concentrations of 50 nM SecA, 2 mM PEP and 0.2 mM NADH when diluted to 100 µl, and incubated at 25 °C for 5 mins. This reaction mixture was added to a 96-well plate with clear, flat-bottomed wells (CoStar). Baseline signal, absorption at 340 nm, was detected for 5 mins. Then 1 µl of 100 mM ATP was added to each well, this was mixed with 5 s double orbital shaking (559 cpm) before measurement for another 30 mins. Then 15 µl of 6.67 µM preprotein was added to reach a final concentration of 1 µM, mixed again, before a continued 30 mins of measurement, resulting first in a background ATPase rate, then SecY-stimulated ATPase rate and finally a preprotein-stimulated ATPase rate phase of the trace (Supplementary Figure S2).

The ATP consumption can be calculated by measuring the gradient of the slope in each phase of the plot (background, ATP, pre-protein), and then rate is calculated using the equation: 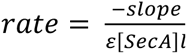. In this equation the extinction coefficient (*ε*) for NADH at 340nM is 6,220 M^-1^.cm^-1^, the concentration of SecA is 50 nM and the path length (*l*) is 0.28 cm. The statistical analysis was performed in GraphPad Prism. For each SecA variant, SecYEG-stimulated and preprotein-stimulated ATPase rates were compared with those of native SecA using multiple unpaired t-tests. Each condition was analysed individually without assuming a consistent standard deviation. The false discovery rate was controlled at Ǫ = 1% using the two-stage step-up method, and comparisons identified as discoveries are marked with an asterisk (*) in Figures 4E and 6E.

### NanoLuc transport assays

The transport experiments were performed using the documented NanoLuc system^48,49^, with split NanoLuc fragments coming together as a result of productive transport, producing a luminescent signal. The large fragment of split NanoLuc (11S) is encapsulated in the proteoliposome lumen, separated from the small split NanoLuc fragment (pep86), which is present in the pre-protein. The proteoliposome membranes are reconstituted with SecYEG, so on the addition of SecA and ATP pre-protein (containing pep86) is translocated across the proteoliposome membrane into the lumen where it encounters 11S, where the fragments rapidly combine to form NanoLuc. The formed NanoLuc then oxidises the substrate Furimazine, present in the reaction mix, to generate a luminescent signal that is detected using a BioTek Synergy Neo2 plate reader. The luminescent signal is proportional to the amount of pep86 (and therefore substrate) transported into the proteoliposome lumen, given excess 11S and Furimazine are present. The reaction master mix is made up in TKM and then furimazine (to a 1/500 – Nano-Glo substrate), Prionex (to 0.125%), 5 mM creatine phosphate, 0.1 mg/ml creatine kinase, it is then incubated at 25 °C for 5 mins before the addition of GST-Dark to 40 µM, and 8.4 nM SecYEG proteoliposomes. Then to each well 1 µM of the relevant SecA and 1 µM pre-protein is added to ensure full saturation of the Sec system.

The NanoLuc experiments begin with an 8 min background signal measurement of the wells, ahead of the addition of 1 mM ATP to each well to initiate transport. Once ATP is added the signal is read for 25 mins. A native SecA trace would display a characteristic lag phase following addition of ATP, followed by a steep (exponential) rise in signal, which eventually plateaus then starts to decay if left long enough. The data processing was performed using GraphPad prism, with the final traces being corrected for background and being fitted, up to the plateau, with the lag exponential model (Supplementary Figure S3).

### Search for other motors

To search for other adapted helicase motors that may contain a domain insert equivalent to PPXD between the N- and C-core of the helicase domain a FoldSeek search was carried out using the 7xha solved structure of SecA. All of the hits from the PDB100 database were then retrieved as FASTA files. A MSA was carried out on these sequences and analysis code was generated to read this MSA and calculate the ratio of residues to dashes between the N- and C-core (dash proportion). The dash proportion was calculated between the equivalent residue positions in the MSA as T216 and F370 in *E. coli* SecA. The results were then filtered by dash proportion to identify the sequences with the highest number of residues between the N- and C-core of the SecA alignment, which could then be inspected to identify if there was a domain insert in this region.

## DECLARATION

The authors declare no competing financial interests.

## DATA AND MATERIALS AVAILABILITY

All data are available in the main text and supplementary information.

## Supplementary Information

### Supplementary Tables

#### Interaction interface from ePISA

PDF of complete output provided – supplementary data 3

#### Co-EV for the co-conserved residues

Table of complete output provided – supplementary data 4

---

**Supplementary figure S1:**
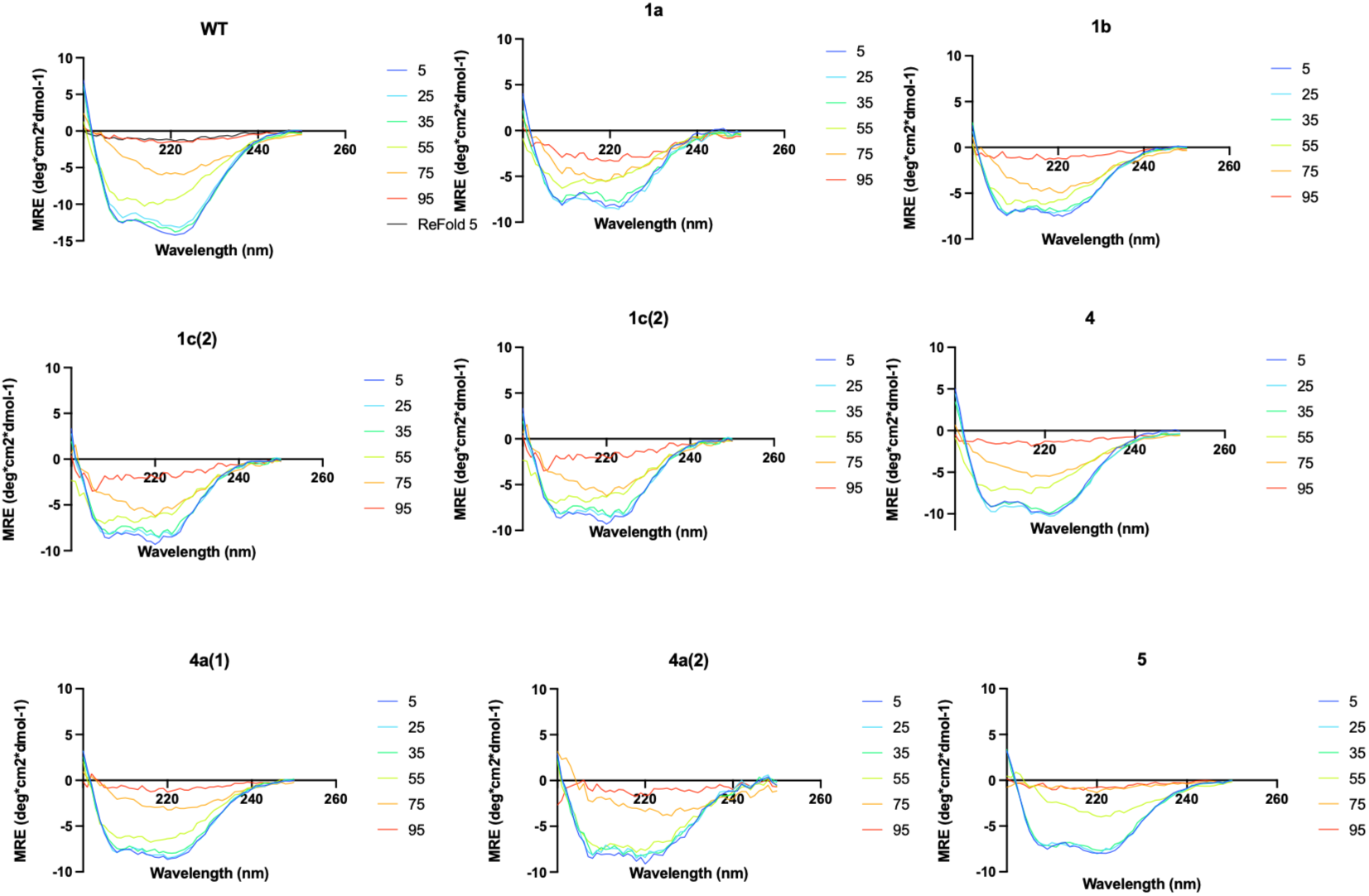
CD spectra of the protein variants used in this study.

**Supplementary figure S2:**
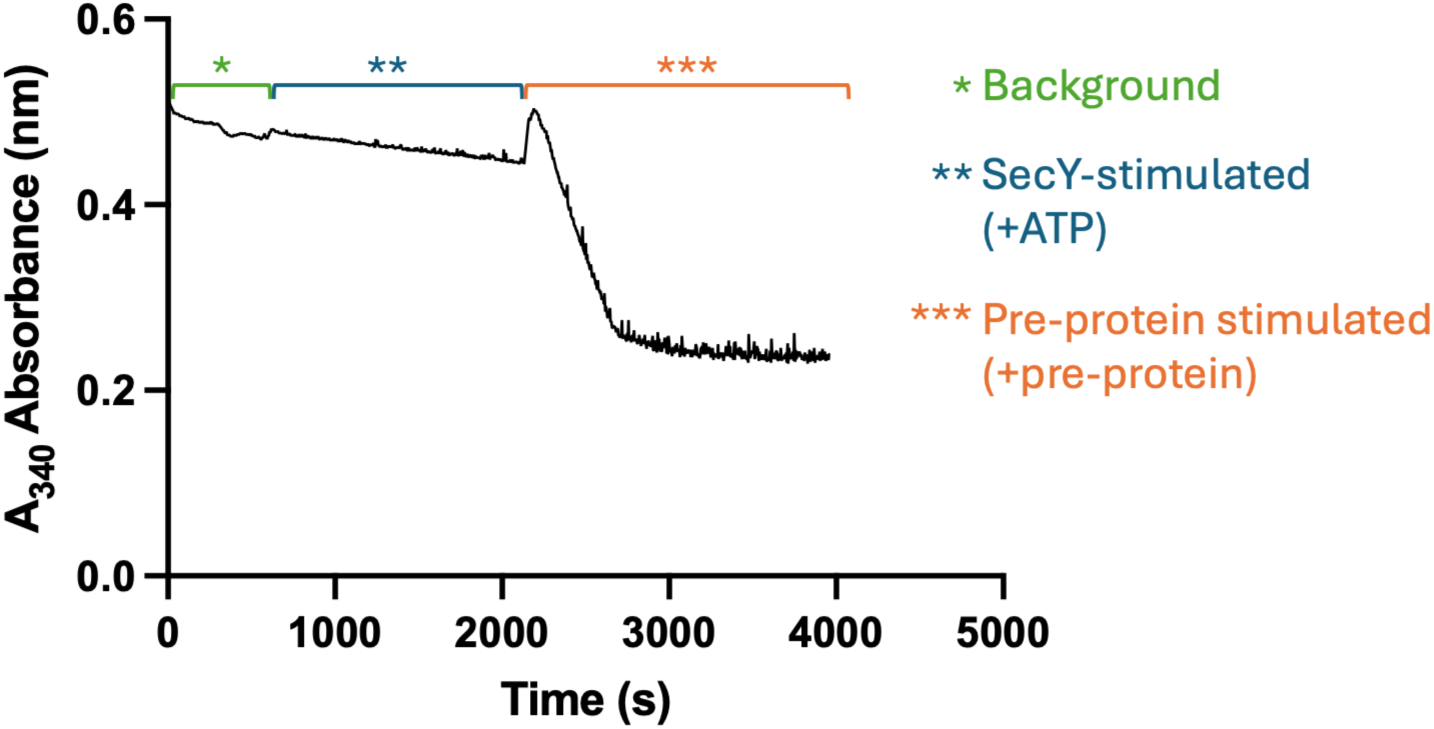
Raw data from the ATPase assay for wild type SecA. The background measurement occurs in the absence of ATP, then ATP is added for the SecY-stimulated rate, followed by the addition of pre-protein to get the pre-protein stimulated rate of SecA ATPase activity.

**Supplementary figure S3:**
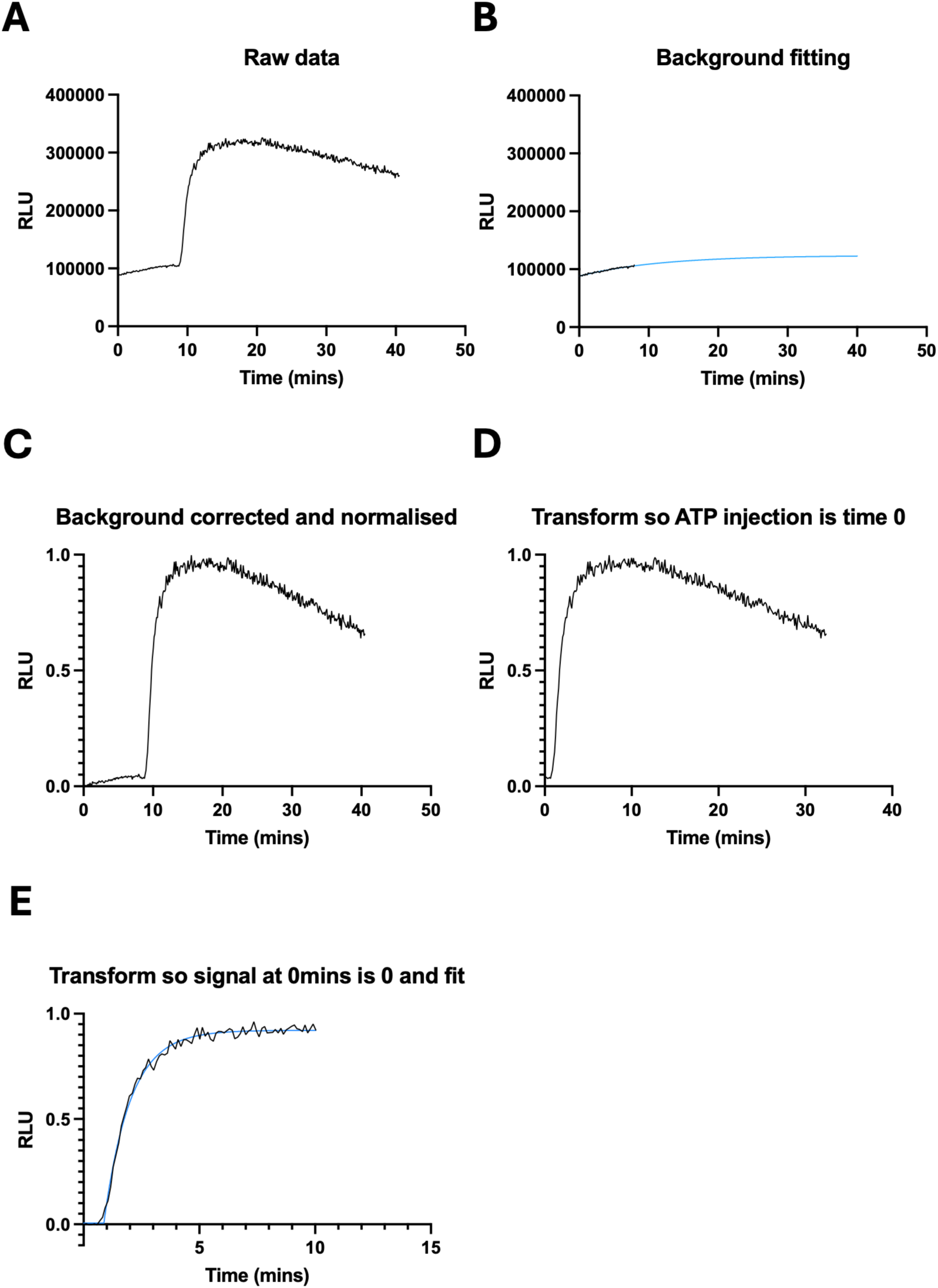
Raw data and the subsequent processing. (A) Raw data for one replicate of wild type SecA. (B) The background signal before the addition of ATP, fitted with a lag exponential with one phase decay and extrapolated for the length of the run. (C) The data having been corrected for the background signal given by the curve from background fitting. This is done for each individual replicate. At this stage the data can be normalized to enable comparison – using a WT reference from each set. (D) The data following the X-transform to begin with the injection of ATP as time point 0. (E) The final data for fitting with the lag exponential with one phase decay having been transformed so there is no RLU signal at time point 0. The data is just plotted up to, and including, the plateau before signal decay. Lag times can be extracted from this analysis, given as X0 from the lag followed by one phase decay curve fit.

**Supplementary table T1:** The top 49 DALI hits using the 7xha B. subtilis structure were other SecA homologues, however beyond that the first 50 hits contained helicases, the top 20 of those hits are shown here.

| Sbjct | z-score | Rmsd | Ali-length | Sbjct-length | Seq-identify | description |
|---|---|---|---|---|---|---|
| 6s0kh | 29.7 | 3.9 | 672 | 838 | 54 | MOLECULE: TRNA-CCA; |
| 6uv3A | 25.8 | 3.4 | 335 | 426 | 13 | MOLECULE: PROBABLE ATP-DEPENDENT RNA HELICASE DDX17; |
| 7pmqD | 25.6 | 3.1 | 322 | 455 | 17 | MOLECULE: ATP-DEPENDENT RNA HELICASE DBPA; |
| 5zc9A | 25.4 | 3.4 | 330 | 378 | 14 | MOLECULE: EUKARYOTIC INITIATION FACTOR 4A-I; |
| 7pmqB | 25.4 | 3.1 | 323 | 457 | 17 | MOLECULE: ATP-DEPENDENT RNA HELICASE DBPA; |
| 6xkiA | 25.3 | 3.5 | 330 | 378 | 14 | MOLECULE: EUKARYOTIC INITIATION FACTOR 4A-I; |
| 7znjf | 25.2 | 3.5 | 334 | 382 | 13 | MOLECULE: EUKARYOTIC INITIATION FACTOR 4A-III, N-TERMINALLY |
| 7pliB | 25.2 | 3.2 | 323 | 456 | 17 | MOLECULE: ATP-DEPENDENT RNA HELICASE DBPA; |
| 6goxA | 25.1 | 7.1 | 585 | 777 | 54 | MOLECULE: PROTEIN TRANSLOCASE SUBUNIT SECA; |
| 7pmqA | 25.1 | 3.2 | 324 | 458 | 17 | MOLECULE: ATP-DEPENDENT RNA HELICASE DBPA; |
| 3fhtB | 25 | 3.4 | 323 | 387 | 15 | MOLECULE: ATP-DEPENDENT RNA HELICASE DDX19B; |
| 7pliE | 25 | 3.3 | 325 | 457 | 17 | MOLECULE: ATP-DEPENDENT RNA HELICASE DBPA; |
| 3fhtA | 24.9 | 3.4 | 329 | 392 | 14 | MOLECULE: ATP-DEPENDENT RNA HELICASE DDX19B; |
| 7pmmB | 24.9 | 3.2 | 325 | 439 | 17 | MOLECULE: ATP-DEPENDENT RNA HELICASE DBPA; |
| 7znjK | 24.8 | 3.5 | 334 | 382 | 13 | MOLECULE: EUKARYOTIC INITIATION FACTOR 4A-III, N-TERMINALLY |
| 7znjF | 24.8 | 3.5 | 335 | 382 | 13 | MOLECULE: EUKARYOTIC INITIATION FACTOR 4A-III, N-TERMINALLY |
| 7znjk | 24.8 | 3.5 | 334 | 382 | 13 | MOLECULE: EUKARYOTIC INITIATION FACTOR 4A-III, N-TERMINALLY |
| 7znja | 24.8 | 3.5 | 334 | 382 | 13 | MOLECULE: EUKARYOTIC INITIATION FACTOR 4A-III, N-TERMINALLY |
| 7znjA | 24.8 | 3.5 | 335 | 382 | 13 | MOLECULE: EUKARYOTIC INITIATION FACTOR 4A-III, N-TERMINALLY |
| 7pliA | 24.7 | 3.2 | 322 | 457 | 17 | MOLECULE: ATP-DEPENDENT RNA HELICASE DBPA; |

**Supplementary table T2:** SecA consensus sequence used in hmmsearch against the full database 700 proteomes dataset. The top 20 hits are shown here.

| full_Evalue | full_score | full_bias | best_dom_Evalue | best_dom_score | best_dom_bias | dom_exp | dom_N | Sequence | Description | Significant |
|---|---|---|---|---|---|---|---|---|---|---|
| 4.90E-308 | 1030.8 | 5.6 | 6.00E-308 | 1030.5 | 5.6 | 1 | 1 | GCA_000154225-GCA_000154225_18450 | Protein translocase subunit SecA | yes |
| 3.20E-121 | 412.6 | 12.8 | 9.90E-120 | 407.6 | 12.8 | 2 | 1 | GCA_003662885-GCA_003662885_12210 | Protein translocase subunit SecA | yes |
| 8.10E-92 | 315.3 | 5 | 3.90E-44 | 157.5 | 0.2 | 3.4 | 3 | GCA_002762795-GCA_002762795_06050 | Protein translocase subunit SecA | yes |
| 0.00096 | 23.8 | 5.9 | 0.00096 | 23.8 | 5.9 | 1.4 | 1 | GCA_001723855-GCA_001723855_21850 | Photosystem I assembly protein Y | yes |
| 0.0011 | 23.6 | 0 | 0.49 | 14.9 | 0 | 2 | 2 | GCA_000194625-GCA_000194625_10230 | putative ATP-dependent RNA helic | yes |
| 0.033 | 18.7 | 1.7 | 0.041 | 18.4 | 1.7 | 1.1 | 1 | GCA_002498125-GCA_002498125_08530 | Chaperone protein DnaJ | no (below inclusion threshold) |
| 0.038 | 18.5 | 0.2 | 0.051 | 18.1 | 0.2 | 1.1 | 1 | GCA_000725425-GCA_000725425_13880 | UvrABC system protein B | no (below inclusion threshold) |
| 0.044 | 18.3 | 0 | 20 | 9.5 | 0 | 2.1 | 2 | GCA_002078355-GCA_002078355_06800 | putative ATP-dependent RNA helic | no (below inclusion threshold) |
| 0.086 | 17.3 | 0 | 0.16 | 16.5 | 0 | 1.3 | 1 | GCA_000213215-GCA_000213215_09870 | putative ATP-dependent RNA helic | no (below inclusion threshold) |
| 0.24 | 15.9 | 0.1 | 0.41 | 15.1 | 0.1 | 1.3 | 1 | GCA_000016525-GCA_000016525_15640 | **ATP-dependent DNA helicase Hel30** | no (below inclusion threshold) |
| 0.25 | 15.8 | 0 | 0.38 | 15.2 | 0 | 1.1 | 1 | GCA_002214545-GCA_002214545_08300 | UvrABC system protein B | no (below inclusion threshold) |
| 0.26 | 15.8 | 0 | 0.59 | 14.6 | 0 | 1.5 | 1 | GCA_000214415-GCA_000214415_16980 | **ATP-dependent DNA helicase Hel30** | no (below inclusion threshold) |
| 0.34 | 15.4 | 0.1 | 0.49 | 14.8 | 0.1 | 1.2 | 1 | GCA_000007305-GCA_000007305_01290 | UvrABC system protein B | no (below inclusion threshold) |
| 0.57 | 14.6 | 0.1 | 0.75 | 14.2 | 0.1 | 1.1 | 1 | GCA_000246985-GCA_000246985_02740 | UvrABC system protein B | no (below inclusion threshold) |
| 1.1 | 13.7 | 0.1 | 1.5 | 13.3 | 0.1 | 1.2 | 1 | GCA_000151105-GCA_000151105_04000 | UvrABC system protein B | no (below inclusion threshold) |
| 1.3 | 13.5 | 0 | 2.2 | 12.7 | 0 | 1.3 | 1 | GCA_000008265-GCA_000008265_14050 | putative ATP-dependent RNA helic | no (below inclusion threshold) |
| 1.3 | 13.4 | 0.1 | 1.7 | 13 | 0.1 | 1.1 | 1 | GCA_000011105-GCA_000011105_01920 | UvrABC system protein B | no (below inclusion threshold) |
| 1.5 | 13.2 | 0 | 1.8 | 13 | 0 | 1 | 1 | GCA_002841105-GCA_002841105_12550 | **ATP-dependent DNA helicase Hel30** | no (below inclusion threshold) |
| 1.5 | 13.2 | 0.9 | 1.7 | 13.1 | 0.9 | 1 | 1 | GCA_002778455-GCA_002778455_07200 | Cell division protein SepF | no (below inclusion threshold) |
| 1.7 | 13 | 0.1 | 4.2 | 11.8 | 0.1 | 1.5 | 1 | GCA_002457555-GCA_002457555_02060 | UvrABC system protein B | no (below inclusion threshold) |

**Supplementary table T3:** SecA consensus sequence used in hmmsearch against the database of compiled helicase sequences from the 700 proteomes dataset. All 6 significant hits are provided here.

| full_Evalue | full_score | full_bias | best_dom_Evalue | best_dom_score | best_dom_bias | dom_exp | dom_N | Sequence | Subfamily | Significant | Evalue_scaled_to_1.76M_db |
|---|---|---|---|---|---|---|---|---|---|---|---|
| 3.5e-06 | 20.3 | 0 | 5.1e-06 | 19.7 | 0 | 1.2 | 1 | GCA_000005845-GCA_000005845_35860 | RecG | yes | 0.011 |
| 2.9e-05 | 17.2 | 0.1 | 5.4e-05 | 16.3 | 0.1 | 1.3 | 1 | GCA_000185805-GCA_000185805_01600 | RecG | yes | 0.094 |
| 7.4e-05 | 15.9 | 0.1 | 0.00013 | 15.1 | 0.1 | 1.3 | 1 | GCA_000016525-GCA_000016525_15640 | RecG | yes | 0.24 |
| 0.00054 | 13 | 0 | 0.00093 | 12.2 | 0 | 1.3 | 1 | GCA_000020945-GCA_000020945_02560 | RecG | yes | 1.7 |
| 0.00074 | 12.6 | 0 | 0.0014 | 11.7 | 0 | 1.3 | 1 | GCA_000724625-GCA_000724625_36190 | Ski2_like | yes | 2.4 |
| 0.0017 | 11.4 | 0.1 | 0.0027 | 10.7 | 0.1 | 1.2 | 1 | GCA_000379665-GCA_000379665_09860 | RigI_like_Y | yes | 5.5 |

**Supplementary table T4:** DALI search hits for SecA PPXD domain from *B. subtilis* (7xha), all the hits with Z-scores above than 8 (significant threshold), and are therefore confident hits, are shown.

| sbjct | z-score | rmsd | ali-length | sbjct-length | seq-identity | description |
|---|---|---|---|---|---|---|
| 6itcA | 23.3 | 0.5 | 136 | 765 | 100 | MOLECULE: PROTEIN TRANSLOCASE SUBUNIT SECA; |
| 8ya0A | 22.5 | 0.4 | 136 | 765 | 100 | MOLECULE: PROTEIN TRANSLOCASE SUBUNIT SECA; |
| 8yasA | 22.5 | 0.5 | 136 | 765 | 100 | MOLECULE: PROTEIN TRANSLOCASE SUBUNIT SECA; |
| 8y9yA | 22.2 | 0.4 | 136 | 765 | 100 | MOLECULE: PROTEIN TRANSLOCASE SUBUNIT SECA; |
| 8y9zA | 21.5 | 0.6 | 136 | 765 | 100 | MOLECULE: PROTEIN TRANSLOCASE SUBUNIT SECA; |
| 9slcA | 21.5 | 1.1 | 135 | 808 | 47 | MOLECULE: PROTEIN TRANSLOCASE SUBUNIT SECA; |
| 7xhaA | 20.5 | 0 | 136 | 765 | 100 | MOLECULE: PROTEIN TRANSLOCASE SUBUNIT SECA; |
| 1tf2A | 20.4 | 3.1 | 132 | 772 | 100 | MOLECULE: PREPROTEIN TRANSLOCASE SECA SUBUNIT; |
| 1tf5A | 20.4 | 2.8 | 130 | 775 | 100 | MOLECULE: PREPROTEIN TRANSLOCASE SECA SUBUNIT; |
| 4ys0A | 19.5 | 2.4 | 134 | 814 | 53 | MOLECULE: PROTEIN TRANSLOCASE SUBUNIT SECA; |
| 7xhbA | 19.2 | 1.4 | 136 | 765 | 93 | MOLECULE: PROTEIN TRANSLOCASE SUBUNIT SECA; |
| 6sxhA | 18.8 | 1.7 | 125 | 769 | 56 | MOLECULE: PROTEIN TRANSLOCASE SUBUNIT SECA 2; |
| 3jv2A | 18.8 | 2.8 | 126 | 758 | 100 | MOLECULE: PROTEIN TRANSLOCASE SUBUNIT SECA; |
| 3jv2B | 18.7 | 3.2 | 131 | 762 | 100 | MOLECULE: PROTEIN TRANSLOCASE SUBUNIT SECA; |
| 3juxA | 17.7 | 2.3 | 131 | 813 | 52 | MOLECULE: PROTEIN TRANSLOCASE SUBUNIT SECA; |
| 6t4hA | 17 | 1.5 | 125 | 771 | 56 | MOLECULE: PROTEIN TRANSLOCASE SUBUNIT SECA 2; |
| 1m6nA | 16.9 | 2.7 | 128 | 802 | 81 | MOLECULE: PREPROTEIN TRANSLOCASE SECA; |
| 5k94B | 16.2 | 2.5 | 129 | 506 | 38 | MOLECULE: MALTOSE-BINDING PERIPLASMIC PROTEIN,PROTEIN TRANS |
| 5k9tA | 16 | 2.7 | 128 | 576 | 43 | MOLECULE: PROTEIN TRANSLOCASE SUBUNIT SECA; |
| 5k94A | 15.7 | 2.5 | 128 | 506 | 41 | MOLECULE: MALTOSE-BINDING PERIPLASMIC PROTEIN,PROTEIN TRANS |
| 1nl3A | 15.3 | 2.3 | 125 | 839 | 39 | MOLECULE: PREPROTEIN TRANSLOCASE SECA 1 SUBUNIT; |
| 3dinB | 14.6 | 1.8 | 129 | 816 | 35 | MOLECULE: PROTEIN TRANSLOCASE SUBUNIT SECA; |
| 1nktB | 14.5 | 2.4 | 126 | 836 | 39 | MOLECULE: PREPROTEIN TRANSLOCASE SECA 1 SUBUNIT; |
| 8ya2A | 14.4 | 0.6 | 136 | 765 | 100 | MOLECULE: PROTEIN TRANSLOCASE SUBUNIT SECA; |
| 2ibmB | 14.4 | 2.4 | 127 | 770 | 85 | MOLECULE: PREPROTEIN TRANSLOCASE SECA SUBUNIT; |
| 3dl8A | 14.2 | 1.8 | 129 | 773 | 57 | MOLECULE: PROTEIN TRANSLOCASE SUBUNIT SECA; |
| 3dl8B | 14.1 | 1.9 | 129 | 773 | 59 | MOLECULE: PROTEIN TRANSLOCASE SUBUNIT SECA; |
| 1nl3B | 13.8 | 2.4 | 125 | 838 | 39 | MOLECULE: PREPROTEIN TRANSLOCASE SECA 1 SUBUNIT; |
| 1nktA | 13.4 | 2.3 | 125 | 836 | 39 | MOLECULE: PREPROTEIN TRANSLOCASE SECA 1 SUBUNIT; |
| 2ipcB | 13.2 | 1.9 | 124 | 934 | 42 | MOLECULE: PREPROTEIN TRANSLOCASE SECA SUBUNIT; |
| 5eulA | 13.1 | 1.1 | 131 | 746 | 88 | MOLECULE: PROTEIN TRANSLOCASE SUBUNIT SECA, INSERTION PEPTI |
| 3dinA | 12.7 | 1.9 | 129 | 816 | 35 | MOLECULE: PROTEIN TRANSLOCASE SUBUNIT SECA; |
| 2vdaA | 12.7 | 3.4 | 130 | 828 | 40 | MOLECULE: TRANSLOCASE SUBUNIT SECA; |
| 2ibmA | 12 | 2.9 | 125 | 780 | 74 | MOLECULE: PREPROTEIN TRANSLOCASE SECA SUBUNIT; |
| 6s0kh | 11.9 | 3.5 | 130 | 838 | 40 | MOLECULE: TRNA-CCA; |
| 3iqmA | 11.4 | 2.3 | 124 | 802 | 81 | MOLECULE: PROTEIN TRANSLOCASE SUBUNIT SECA; |
| 2ipcC | 11.1 | 2.3 | 128 | 939 | 43 | MOLECULE: PREPROTEIN TRANSLOCASE SECA SUBUNIT; |
| 3iqyA | 11 | 2.2 | 124 | 802 | 83 | MOLECULE: PROTEIN TRANSLOCASE SUBUNIT SECA; |
| 1m74A | 10.9 | 2 | 125 | 802 | 76 | MOLECULE: PREPROTEIN TRANSLOCASE SECA; |
| 2ipcA | 10.9 | 2.4 | 128 | 939 | 41 | MOLECULE: PREPROTEIN TRANSLOCASE SECA SUBUNIT; |
| 2ipcD | 10.8 | 2 | 125 | 934 | 42 | MOLECULE: PREPROTEIN TRANSLOCASE SECA SUBUNIT; |
| 4uaqA | 10.4 | 2.5 | 113 | 705 | 33 | MOLECULE: PROTEIN TRANSLOCASE SUBUNIT SECA 2; |
| 6goxA | 7.1 | 3.1 | 98 | 777 | 44 | MOLECULE: PROTEIN TRANSLOCASE SUBUNIT SECA; |

**Supplementary table T5:** A table of the hits from the B. subtilis PPXD domain using blastp with the BLOSUM62 matrix with default gap cost, and a very permissive expected value of 100 to try and identify distant homologues. Hitlist size was set to 100 but only 29 sequences were returned, of which only the final 3 were not SecA homologues and are likely to be noise with E-values above 1.

| Description | Scientific Name | Max Score | Total Score | Query Cover | E value | Per. ident | Acc. Len | Accession |
|---|---|---|---|---|---|---|---|---|
| Chain A, Protein translocase subunit SecA [Bacillus subtilis subsp. subtilis str. 168] | Bacillus subtilis subsp. subtilis str. 168 | 279 | 279 | 100% | 2.00E-90 | 100 | 765 | 8YA0_A |
| Chain A, Preprotein translocase secA subunit [Bacillus subtilis] | Bacillus subtilis | 278 | 278 | 100% | 4.00E-90 | 100 | 780 | 2IBM_A |
| Chain A, Protein translocase subunit secA [Bacillus subtilis] | Bacillus subtilis | 278 | 278 | 100% | 4.00E-90 | 100 | 779 | 3DL8_A |
| Chain A, Protein translocase subunit SecA [Bacillus subtilis subsp. subtilis str. 168] | Bacillus subtilis subsp. subtilis str. 168 | 278 | 278 | 100% | 4.00E-90 | 100 | 778 | 7XHA_A |
| Chain A, Protein translocase subunit SecA [Bacillus subtilis] | Bacillus subtilis | 278 | 278 | 100% | 5.00E-90 | 100 | 783 | 3JV2_A |
| Chain A, Protein translocase subunit secA [Bacillus subtilis subsp. subtilis str. 168] | Bacillus subtilis subsp. subtilis str. 168 | 278 | 278 | 100% | 5.00E-90 | 100 | 802 | 3IQM_A |
| Chain A, Preprotein translocase secA [Bacillus subtilis] | Bacillus subtilis | 278 | 278 | 100% | 5.00E-90 | 100 | 802 | 1M6N_A |
| Chain A, Protein translocase subunit secA [Bacillus subtilis] | Bacillus subtilis | 278 | 278 | 100% | 1.00E-89 | 100 | 841 | 3IQY_A |
| Chain A, Protein translocase subunit SecA, Insertion Peptide Chimera [synthetic construct] | synthetic construct | 278 | 278 | 100% | 1.00E-89 | 100 | 836 | 5EUL_A |
| Chain A, Preprotein translocase secA subunit [Bacillus subtilis] | Bacillus subtilis | 278 | 278 | 100% | 2.00E-89 | 100 | 844 | 1TF2_A |
| Chain A, Protein translocase subunit SecA 2 [Clostridioides difficile 630] | Clostridioides difficile 630 | 171 | 171 | 100% | 5.00E-50 | 59.56 | 786 | 6SXH_A |
| Chain A, Protein translocase subunit SecA 2 [Clostridioides difficile 630] | Clostridioides difficile 630 | 171 | 171 | 100% | 5.00E-50 | 59.56 | 781 | 6T4H_A |
| Chain A, Protein translocase subunit SecA [Thermotoga maritima MSB8] | Thermotoga maritima MSB8 | 147 | 147 | 100% | 2.00E-41 | 53.68 | 824 | 4YS0_A |
| Chain A, Protein translocase subunit secA [Thermotoga maritima] | Thermotoga maritima | 147 | 147 | 100% | 2.00E-41 | 53.68 | 822 | 3JUX_A |
| Chain A, Protein translocase subunit secA [Thermotoga maritima MSB8] | Thermotoga maritima MSB8 | 147 | 147 | 100% | 2.00E-41 | 53.68 | 871 | 3DIN_A |
| Chain A, Preprotein translocase SecA subunit [Thermus thermophilus HB8] | Thermus thermophilus HB8 | 145 | 145 | 100% | 1.00E-40 | 52.03 | 997 | 2IPC_A |
| Chain A, Protein translocase subunit SecA [Escherichia coli 55989] | Escherichia coli 55989 | 131 | 131 | 100% | 5.00E-36 | 43.51 | 578 | 5K9T_A |
| Chain A, Preprotein translocase secA subunit [Escherichia coli] | Escherichia coli | 131 | 131 | 100% | 9.00E-36 | 43.51 | 853 | 2FSF_A |
| Chain h, Protein translocase subunit SecA [Escherichia coli] | Escherichia coli | 131 | 131 | 100% | 9.00E-36 | 43.51 | 838 | 6S0K_h |
| Chain A, Protein translocase subunit SecA [Escherichia coli K-12] | Escherichia coli K-12 | 131 | 131 | 100% | 9.00E-36 | 43.51 | 822 | 6GOX_A |
| Chain A, TRANSLOCASE SUBUNIT SECA [Escherichia coli] | Escherichia coli | 131 | 131 | 100% | 9.00E-36 | 43.51 | 828 | 2VDA_A |
| Chain A, Preprotein translocase secA subunit [Escherichia coli] | Escherichia coli | 128 | 128 | 100% | 1.00E-34 | 42.86 | 853 | 2FSG_A |
| Chain A, Preprotein translocase secA 1 subunit [Mycobacterium tuberculosis] | Mycobacterium tuberculosis | 124 | 124 | 100% | 3.00E-33 | 44.85 | 922 | 1NKT_A |
| Chain A, Protein translocase subunit SecA [Mycoplasmoides pneumoniae M129] | Mycoplasmoides pneumoniae M129 | 121 | 121 | 100% | 2.00E-32 | 47.79 | 808 | 9SLC_A |
| Chain A, Maltose-binding periplasmic protein,Protein translocase subunit SecA,Maltose-binding periplasmic protein [Escherichia coli O157:H7] | Escherichia coli O157:H7 | 110 | 110 | 90% | 1.00E-28 | 40.71 | 508 | 5K94_A |
| Chain A, Protein translocase subunit SecA 2 [Mycobacterium tuberculosis H37Rv] | Mycobacterium tuberculosis H37Rv | 79.7 | 79.7 | 100% | 1.00E-17 | 35.46 | 778 | 4UAQ_A |
| Chain h3, Coiled-coil domain-containing protein 40 [Homo sapiens] | Homo sapiens | 28.9 | 28.9 | 36% | 4.4 | 36.73 | 1142 | 8J07_h3 |
| Chain Dz, Coiled-coil domain containing 40 [Bos taurus] | Bos taurus | 27.7 | 27.7 | 32% | 12 | 36.36 | 1125 | 9FQR_Dz |
| Chain F, PAP3 [Sinapis alba] | Sinapis alba | 25.4 | 25.4 | 26% | 75 | 27.78 | 675 | 8QMA_F |

**Supplementary Table T6:**
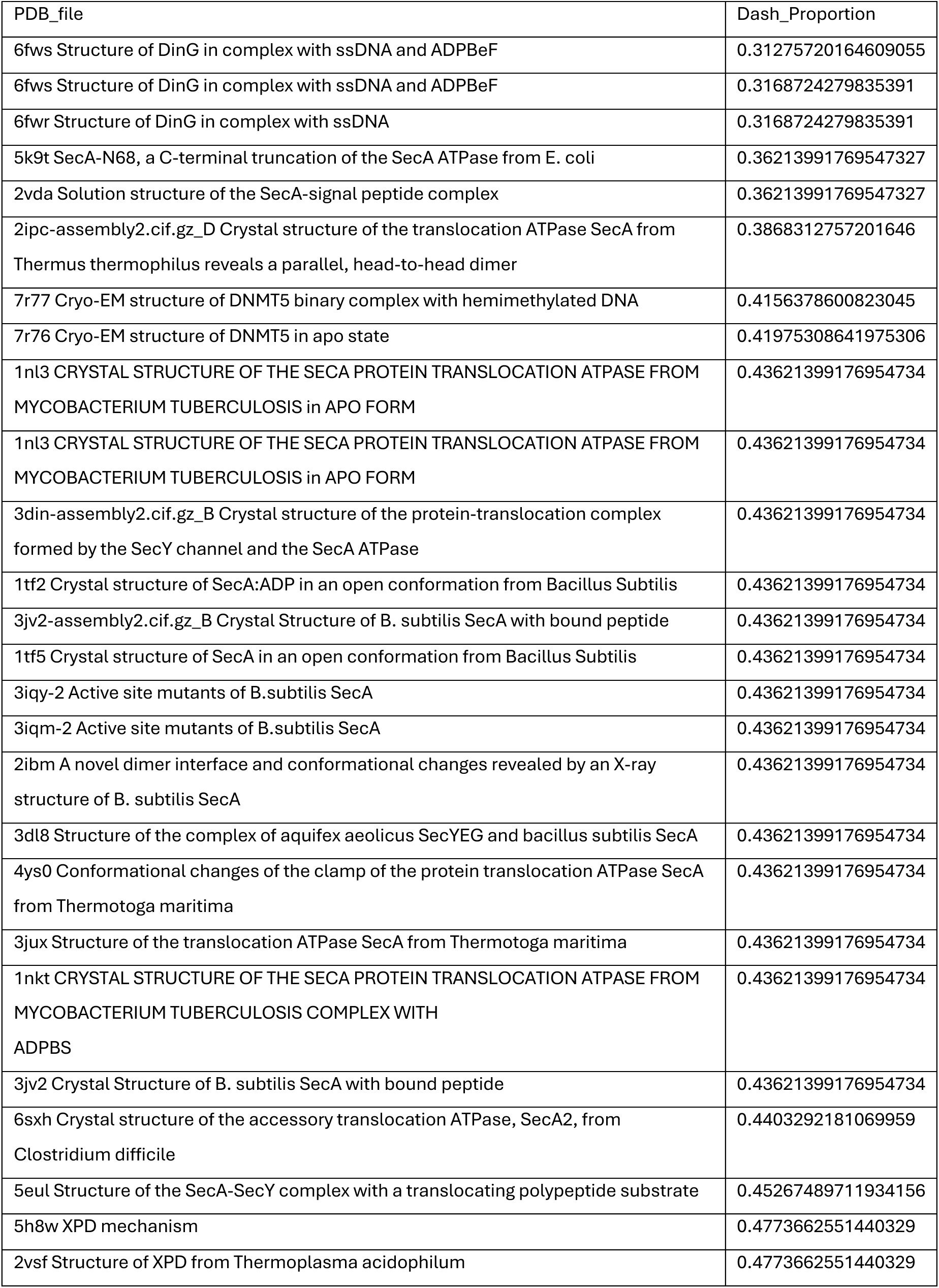
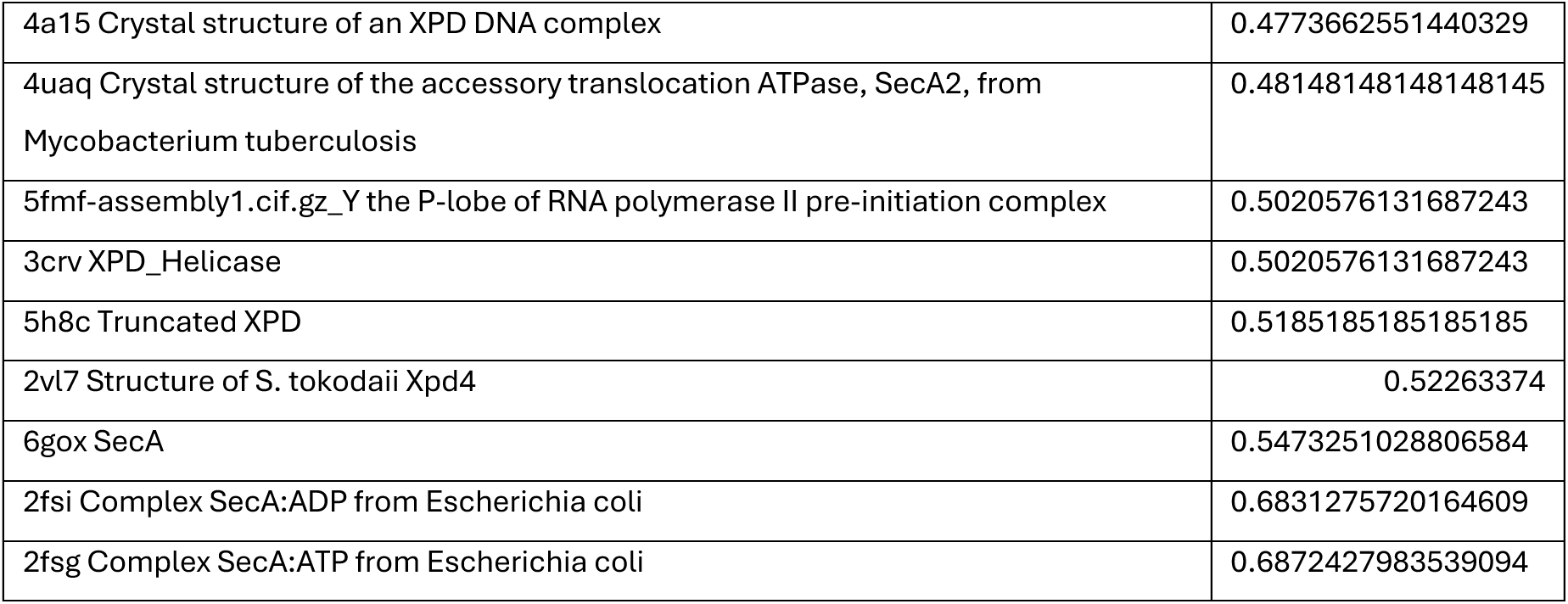
The only proteins with a potential domain insert between N-core and C-core when investigating the SecA FoldSeek results are SecA proteins from other organisms and XPD helicases which do handle nucleic acids. All the hits from the search are provided.

| PDB_file | Dash_Proportion |
|---|---|
| 6fws Structure of DinG in complex with ssDNA and ADPBeF | 0.31275720164609055 |
| 6fws Structure of DinG in complex with ssDNA and ADPBeF | 0.3168724279835391 |
| 6fwr Structure of DinG in complex with ssDNA | 0.3168724279835391 |
| 5k9t SecA-N68, a C-terminal truncation of the SecA ATPase from E. coli | 0.36213991769547327 |
| 2vda Solution structure of the SecA-signal peptide complex | 0.36213991769547327 |
| 2ipc-assembly2.cif.gz_D Crystal structure of the translocation ATPase SecA from Thermus thermophilus reveals a parallel, head-to-head dimer | 0.3868312757201646 |
| 7r77 Cryo-EM structure of DNMT5 binary complex with hemimethylated DNA | 0.4156378600823045 |
| 7r76 Cryo-EM structure of DNMT5 in apo state | 0.41975308641975306 |
| 1nl3 CRYSTAL STRUCTURE OF THE SECA PROTEIN TRANSLOCATION ATPASE FROM MYCOBACTERIUM TUBERCULOSIS in APO FORM | 0.43621399176954734 |
| 1nl3 CRYSTAL STRUCTURE OF THE SECA PROTEIN TRANSLOCATION ATPASE FROM MYCOBACTERIUM TUBERCULOSIS in APO FORM | 0.43621399176954734 |
| 3din-assembly2.cif.gz_B Crystal structure of the protein-translocation complex formed by the SecY channel and the SecA ATPase | 0.43621399176954734 |
| 1tf2 Crystal structure of SecA:ADP in an open conformation from Bacillus Subtilis | 0.43621399176954734 |
| 3jv2-assembly2.cif.gz_B Crystal Structure of B. subtilis SecA with bound peptide | 0.43621399176954734 |
| 1tf5 Crystal structure of SecA in an open conformation from Bacillus Subtilis | 0.43621399176954734 |
| 3iqy-2 Active site mutants of B.subtilis SecA | 0.43621399176954734 |
| 3iqm-2 Active site mutants of B.subtilis SecA | 0.43621399176954734 |
| 2ibm A novel dimer interface and conformational changes revealed by an X-ray structure of B. subtilis SecA | 0.43621399176954734 |
| 3dl8 Structure of the complex of aquifex aeolicus SecYEG and bacillus subtilis SecA | 0.43621399176954734 |
| 4ys0 Conformational changes of the clamp of the protein translocation ATPase SecA from Thermotoga maritima | 0.43621399176954734 |
| 3jux Structure of the translocation ATPase SecA from Thermotoga maritima | 0.43621399176954734 |
| 1nkt CRYSTAL STRUCTURE OF THE SECA PROTEIN TRANSLOCATION ATPASE FROM MYCOBACTERIUM TUBERCULOSIS COMPLEX WITH<br>ADPBS | 0.43621399176954734 |
| 3jv2 Crystal Structure of B. subtilis SecA with bound peptide | 0.43621399176954734 |
| 6sxh Crystal structure of the accessory translocation ATPase, SecA2, from Clostridium difficile | 0.4403292181069959 |
| 5eul Structure of the SecA-SecY complex with a translocating polypeptide substrate | 0.45267489711934156 |
| 5h8w XPD mechanism | 0.4773662551440329 |
| 2vsf Structure of XPD from Thermoplasma acidophilum | 0.4773662551440329 |
| 4a15 Crystal structure of an XPD DNA complex | 0.4773662551440329 |
| 4uaq Crystal structure of the accessory translocation ATPase, SecA2, from Mycobacterium tuberculosis | 0.48148148148148145 |
| 5fmf-assembly1.cif.gz_Y the P-lobe of RNA polymerase II pre-initiation complex | 0.5020576131687243 |
| 3crv XPD_Helicase | 0.5020576131687243 |
| 5h8c Truncated XPD | 0.5185185185185185 |
| 2vl7 Structure of S. tokodaii Xpd4 | 0.52263374 |
| 6gox SecA | 0.5473251028806584 |
| 2fsi Complex SecA:ADP from Escherichia coli | 0.6831275720164609 |
| 2fsg Complex SecA:ATP from Escherichia coli | 0.6872427983539094 |

